# A network of interacting ciliary tip proteins with opposing activities imparts slow and processive microtubule growth

**DOI:** 10.1101/2024.03.25.586532

**Authors:** Harriet A. J. Saunders, Cyntha M. van den Berg, Robin Hoogebeen, Donna Schweizer, Kelly E. Stecker, Ronald Roepman, Stuart C. Howes, Anna Akhmanova

## Abstract

Cilia are essential motile or sensory organelles found on many eukaryotic cells. Their formation and function rely on axonemal microtubules, which exhibit very slow dynamics, however the underlying biochemical mechanisms are largely unexplored. Here, we reconstituted *in vitro* the individual and collective activities of the ciliary tip module proteins, CEP104, CSPP1, TOGARAM1, ARMC9 and CCDC66, which interact with each other and with microtubules, and, when mutated, cause ciliopathies such as Joubert syndrome. CEP104, a protein containing a tubulin-binding TOG domain, is an inhibitor of microtubule growth and shortening that interacts with EBs on the microtubule surface and with a luminal microtubule-pausing factor CSPP1. Another TOG-domain protein, TOGARAM1, overcomes growth inhibition imposed by CEP104 and CSPP1. CCDC66 and ARMC9 do not affect microtubule dynamics directly but act as scaffolds for their partners. Cryo-electron tomography showed that together, ciliary tip module members form plus-end-specific cork-like structures which reduce protofilament flaring. The combined effect of these proteins is very slow processive microtubule elongation, which recapitulates axonemal dynamics in cells.

## Introduction

Cilia are motile or sensory organelles present on the surface of many eukaryotic cells. Motile cilia generate movement for processes such as mucus flow in airways or swimming of unicellular green algae, while primary cilia function as signaling hubs with fundamental roles in organism development (Klena and Pigino, 2022; Mill et al., 2023). Ciliary dysfunction is associated with numerous diseases, collectively called ciliopathies, which affect multiple organs and tissues (Badano et al., 2006; Hildebrandt et al., 2011; Klena and Pigino, 2022). The core of each cilium is formed by a microtubule (MT)-based structure, the axoneme, which is often disrupted in ciliopathies (Deretic et al., 2023). Diverse proteins and pathways responsible for axoneme formation have been identified through genetic screens and cell biology studies, yet the biochemical mechanisms governing the dynamics of axonemal MTs remain very poorly understood (Keeling et al., 2016).

Unlike cytoplasmic MTs, axonemal plus ends grow very slowly likely impeding the formation of a stabilizing GTP cap, which is required for preventing MT disassembly (Desai and Mitchison, 1997) and efficient tubulin addition (Wieczorek et al., 2015). The absence of a stable GTP cap can be compensated by specific factors bound to MT ends. Excellent candidates for this function in cilia are the components of the recently identified ciliary tip interaction network or “module”, CEP104, CSPP1, TOGARAM1, ARMC9 and CCDC66 (Latour et al., 2020). Depletion of any of these proteins causes ciliary defects, and mutations in most of them lead to a primary ciliopathy called Joubert syndrome, characterized by brain malformations, breathing problems or more severe developmental disorders (Akizu et al., 2014; Hildebrandt et al., 2011; Latour et al., 2020; Shaheen et al., 2014; Srour et al., 2015; Tuz et al., 2014; Van De Weghe et al., 2017). The proteins comprising the ciliary tip module interact with MTs and with each other, and although they do not form a stoichiometric complex, they are all important for controlling ciliary length and the signaling pathways dependent on primary cilia (Conkar et al., 2017; Das et al., 2015; Frikstad et al., 2019; Jiang et al., 2012; Latour et al., 2020; Odabasi et al., 2023; Patzke et al., 2006; Van De Weghe et al., 2017). In addition to their roles in cilia, some of the tip module proteins also associate with centrioles, centrosomes, centriolar satellites and cytoplasmic MTs, where they participate in cilia-independent processes, such as cell division (Batman et al., 2022; Conkar et al., 2019; Patzke et al., 2006; Ryniawec et al., 2023; Zhu et al., 2015).

CEP104 (also known as FAP256), TOGARAM1 (also known as Crescerin or CHE-12) and ARMC9 are conserved across eukaryotes, including *Chlamydomonas*, *Tetrahymena* and *C. elegans*, where they participate in the biogenesis of motile or sensory cilia (Bacaj et al., 2008; Das et al., 2015; Louka et al., 2018; Perlaza et al., 2022; Satish Tammana et al., 2013). Both CEP104 and TOGARAM1 have evolutionary conserved TOG domains (Fig. 1A). TOGARAM1 contains four TOG domains, two of which can promote tubulin polymerization in light scattering experiments (Das et al., 2015), and the same is true for the single tubulin-binding TOG domain of CEP104 (Al-Jassar et al., 2017; Rezabkova et al., 2016; Yamazoe et al., 2020). Loss of function mutations in TOGARAM1 and CEP104 in multiple organisms result in short cilia (Bacaj et al., 2008; Das et al., 2015; Frikstad et al., 2019; Latour et al., 2020; Louka et al., 2018; Perlaza et al., 2022; Satish Tammana et al., 2013; Yamazoe et al., 2020). Similarly, loss of CSPP1, a protein that increases MT stability by binding inside the MT lumen and promoting pausing (van den Berg et al., 2023; Wang et al., 2024), results in short cilia (Frikstad et al., 2019). ARMC9 on its own cannot bind to MTs but acts as a scaffold for other ciliary tip module components (Latour et al., 2020). ARMC9 and TOGARAM1 have been localized to motile cilia, where they were found to have opposite functions in regulating MT length (Louka et al., 2018; Perlaza et al., 2022), though mutants of both proteins make mammalian primary cilia shorter (Latour et al., 2020). Finally, CSPP1 and CEP104 interact with CCDC66, and co-depletion of CCDC66 with either CSPP1 or CEP104 results in further reduction of ciliary length compared to the removal of either CSPP1 or CEP104 alone (Odabasi et al., 2023).

**Figure 1.**
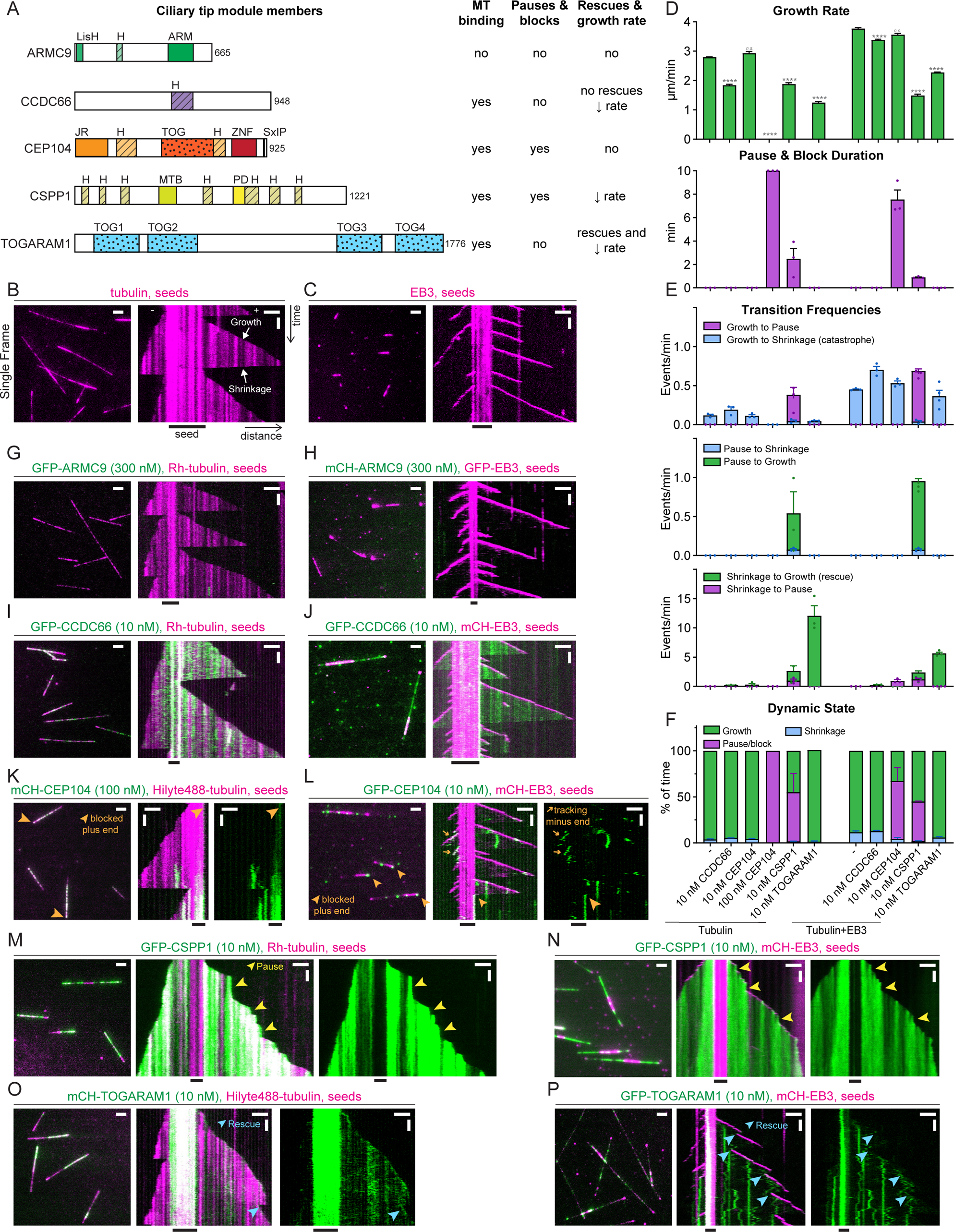
Ciliary tip module proteins have distinct effects on MT dynamics *in vitro*. (A) Schematic representation of ciliary tip module members and summary table highlighting individual protein effects on MT dynamics. Jelly-roll, JR; zinc finger, ZNF; MT binding, MTB; pause domain, PD; α-helical domain, H. (B-C) Fields of view (left, scale bar 2 µm) and kymographs (right, scale bars 2 µm and 60 s) illustrating MT dynamics from GMPCPP-stabilized seeds with either 15 µM tubulin supplemented with 3% HiLyte488-labelled tubulin or 20 nM GFP-EB3. (D-F) Parameters of MT plus-end dynamics in the presence of 15 µM tubulin alone or with 20 nM EB3 in combination with indicated concentrations of proteins (from kymographs shown in B-C, I-P, and S1C). (D) Bars represent pooled data from three independent experiments (growth rate) or averaged means from three independent experiments (pause and block duration), total number of growth events, pauses or blocks: tubulin alone, n=356, 0; 10 nM CCDC66, n=411, 0; 10 nM CEP104, n=306, 0; 100 nM CEP104, n=0, 138; 10 nM CSPP1,n=422, 97; 10 nM TOGARAM1, n=347, 0; EB3 alone, n=938, 0; EB3 with 10 nM CCDC66, n=562, 0; EB3 with 10 nM CEP104, n=213, 101; EB3 with CSPP1, n=1715, 273; EB3 with TOGARAM1, n=861, 0. ****, p<0.0001; n.s., not significant; Kruskal-Wallis test followed by Dunńs post-test, all conditions were compared to their relevant control (either tubulin alone or tubulin with 20 nM EB3). (E-F) Bars represent averaged means from three independent experiments. Error bars represent s.e.m.. Movies were acquired for 10 minutes, therefore this is the maximum time for pause duration, (G-P) Fields of view (left, scale bar 2 µm) and kymographs (right, scale bars 2 µm and 60 s) illustrating MT dynamics from GMPCPP-stabilized seeds with either 15 µM tubulin supplemented with 3% HiLyte488 or Rhodamine-labelled tubulin, or 20 nM GFP- or mCherry-EB3, and indicated concentrations and colors of ciliary tip module proteins. Orange arrowheads, blocked plus ends; orange arrows, CEP104 tracking minus ends; yellow arrowheads, pauses; blue arrowheads, rescues.

To uncover biochemical mechanisms underlying the function of these proteins and their combined regulation of ciliary MTs, we reconstituted *in vitro* their individual and collective effects on MT dynamics. Although the individual TOG domain of CEP104 was previously shown to promote MT polymerization (Yamazoe et al., 2020), we found that the full-length protein specifically inhibited MT plus-end elongation and shortening. This activity was potentiated by EB3, CSPP1 and CCDC66, which could all recruit CEP104 to MTs. This was surprising as EB3 and CSPP1 bind to the outer and inner surface of the MT wall, respectively (Maurer et al., 2012; van den Berg et al., 2023). TOGARAM1 functioned as a polymerase that converted the inhibition of MT polymerization imposed by CEP104 into slow growth. ARMC9 and CCDC66 behaved as scaffolds, enhancing the effects of their binding partners. The combination of all five ciliary tip module proteins resulted in very slow and processive MT plus-end elongation, an effect that required TOGARAM1 and became less robust when one of the other proteins was left out. MT growth rates in these conditions were in the same range as initial elongation rates of regenerating flagella of single-celled organisms (Rosenbaum and Child, 1967; Rosenbaum et al., 1969; Witman, 1975). Cryo-electron tomography showed that together, ciliary tip proteins formed a champagne cork-like structure that protruded from MT plus ends and diminished protofilament flaring. Altogether, our findings demonstrate that, through a combination of opposing activities, ciliary tip module components can stabilize MT plus ends and drive their slow but persistent elongation.

## Results

### Ciliary tip module proteins have distinct effects on MT dynamics in vitro

To characterize the effects of ciliary tip module proteins on MT polymerization, we N-terminally tagged them with mCherry or GFP and purified them from transiently transfected HEK293 cells (Fig. 1A, S1A). Mass spectrometry-based analysis demonstrated that the protein preparations contained only very minor contaminations with other known regulators of MT dynamics, such as CSPP1 (Fig. S1B). In addition, the heat shock protein Hsp70, a protein that to our knowledge has no effect on MT dynamics, was present in the preparations of CCDC66 and CSPP1 (Fig. S1B). We used these proteins to perform *in vitro* reconstitution assays with MTs grown from GMPCPP-stabilized seeds and observed their impact on MT dynamics by Total Internal Reflection Fluorescence (TIRF) microscopy (Bieling et al., 2007; van den Berg et al., 2023). Although axonemes are composed of MT doublets, single MTs are appropriate substrates to study the effects of ciliary tip proteins, because doublets become singlets at ciliary tips (Fisch and Dupuis-Williams, 2011; Kiesel et al., 2020; Legal et al., 2023). To detect MTs, we used either fluorescent tubulin or fluorescently tagged EB3, which increases MT growth rate and induces catastrophes (Komarova et al., 2009) (Fig. 1B-F). Studying the effects of ciliary tip proteins in the presence of EBs is relevant, because EB1 and EB3 localize to axonemal MTs in both primary and motile cilia (Kiesel et al., 2020; Pedersen et al., 2003; Schroder et al., 2007).

Among the tested proteins, ARMC9 was the only one that displayed no MT binding even when present at 300 nM (Fig. 1G and H), in agreement with data in cells (Latour et al., 2020). CCDC66, which is known to interact with MTs (Batman et al., 2022), bound along the MT lattice already at 10 nM and slightly decreased the growth rate, but did not affect the frequencies of catastrophes and rescues and did not induce pausing (Fig. 1D-F, I and J).

CEP104 could not autonomously bind MTs at 10 nM in assays with tubulin alone (Fig. S1C). However, at 100 nM it efficiently blocked MT growth specifically at the plus end (Fig. 1D-F and K), as confirmed by including in the assay a constitutively active fragment of the plus-end directed Kinesin-1 DmKHC(1-421) (Fig. S1D). In the presence of its binding partner EB3 (Jiang et al., 2012), CEP104 could specifically inhibit MT plus-end polymerization already at 10 nM (Fig. 1D-F and L). CEP104 colocalized with EB3 at some growing plus and occasionally also minus ends, however, this did not alter growth rates at the plus end (Fig.1D and L). Once MT growth was arrested, no regrowth was detected for the remainder of the experiment (Fig 1E, K, and L).

Episodes of inhibited growth, albeit transient ones, were also observed with CSPP1, as we have shown previously (Fig. 1D-F, M and N) (van den Berg et al., 2023). Both with and without EB3 CSPP1 preferentially binds at both plus and minus ends when they undergo growth perturbations (van den Berg et al., 2023; Wang et al., 2024), and therefore it displayed discreet sites of increased lattice accumulation (Fig. 1M and N). At the plus ends, CSPP1 accumulations decreased growth rate, induced pauses and facilitated transitions from pause to growth, whereas pausing was less obvious at the minus ends due to their overall slower dynamics (Fig. 1D-F, M and N).

Finally, TOGARAM1 bound to MT plus ends, reduced their growth rate and induced transitions from shrinkage to growth (rescues), and these effects were not altered by the presence of EB3 (Fig. 1D-F, O and P). A decrease in MT growth rate was unexpected, as TOGARAM1 structurally resembles the MT polymerase ch-TOG/XMAP215 (Das et al., 2015), which strongly accelerates MT polymerization (Brouhard et al., 2008). Altogether, we show that all the full-length members of the ciliary tip module can be purified and used for *in vitro* assays, where they display opposing effects on MT dynamics.

### Plus-end blocking by CEP104 depends on the TOG domain and is potentiated by EB3, CCDC66 and CSPP1

Among the ciliary tip proteins analyzed above, the effect of CEP104 was the most unexpected as it was thought to be a MT polymerase (Yamazoe et al., 2020), and we therefore set out to dissect it in more detail. Single molecule counting experiments demonstrated that CEP104 forms a dimer, in agreement with the sedimentation profile of its first helical domain (Fig. 2A) (Rezabkova et al., 2016). In the presence of EB3, ∼5-6 CEP104 molecules (2-3 dimers) were sufficient to block MT plus-end growth (Fig. 2B). In these conditions, Fluorescence Recovery after Photobleaching (FRAP) showed that CEP104 did not exchange at blocked MT plus ends (Fig. 2C), whereas EB3, associated with the same ends, did partially turn over, albeit significantly slower than on growing MT plus ends (Fig. 2D and E). This could be explained by the direct interaction of EB3 and CEP104 on the MT through the SxIP motif of CEP104, SKIP (Fig. 3A) (Jiang et al., 2012). Mutation of this motif has previously been shown to abolish the interactions between the C-terminus of EB and its partners (Honnappa et al., 2009; Yamazoe et al., 2020). Indeed, CEP104 SKNN double mutation prevented EB3-dependent recruitment to MTs; therefore, even in the presence of EB3, 100 nM CEP104 SKNN was needed to block plus-end growth (Fig. 3A-D, and S2A).

**Figure 2.**
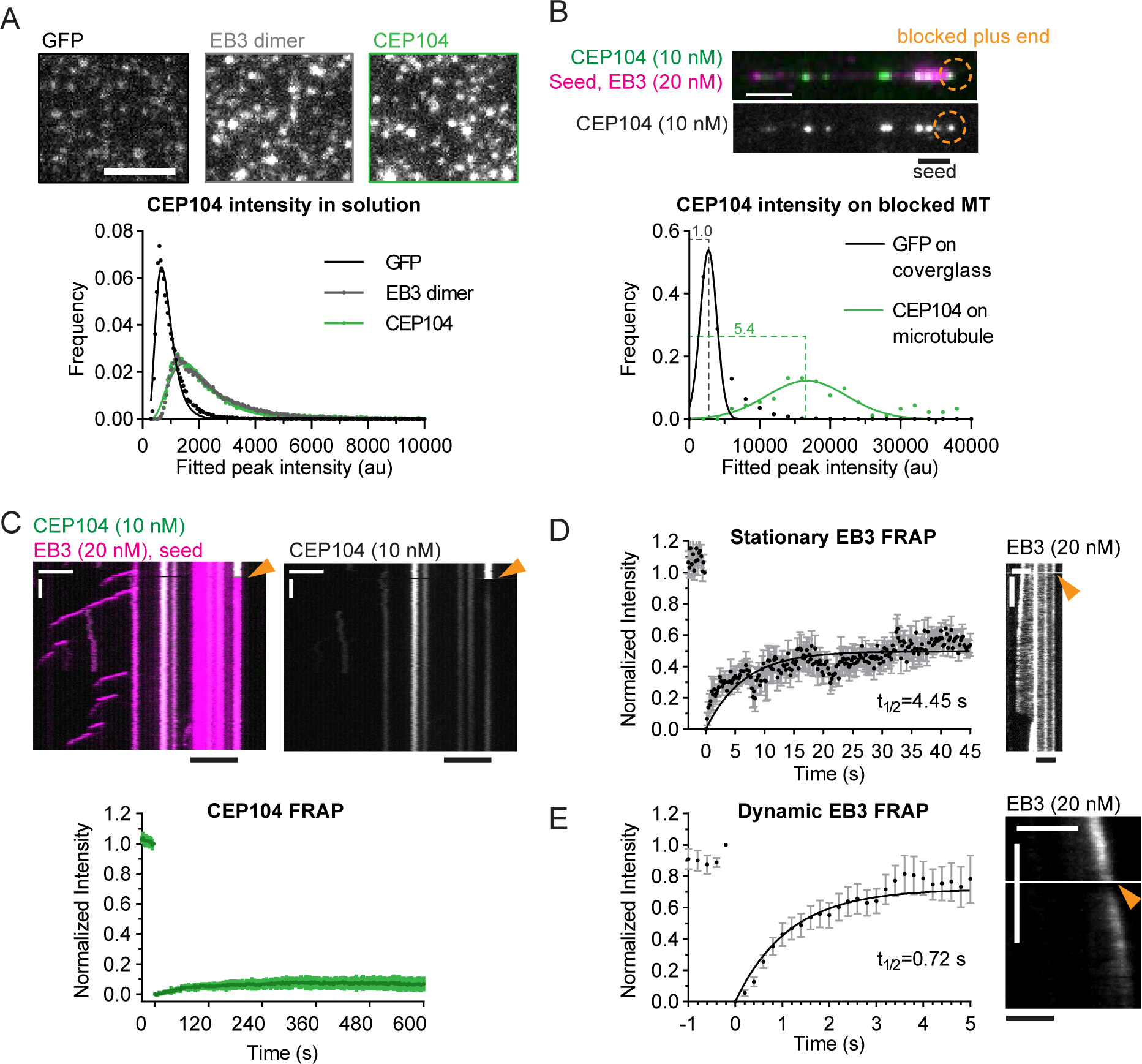
A few CEP104 molecules stably block MT plus ends. (A) Fields of view (top) and histogram plot (bottom) of fluorescent intensities of single GFP molecules, GFP-EB3 dimers, and GFP-CEP104 dimers immobilized in separate chambers of the same coverslip. Number of molecules analyzed: GFP, n=29981; GFP-EB3, n=55378; GFP-CEP104, n=32335. Scale bar 2 µm. (B) Representative MT with CEP104-blocked plus end (top) and histogram plot (bottom) of fluorescent intensities of single GFP molecules and GFP-CEP104 intensity at blocked plus end immobilized in separate chambers of the same coverslip. Number of molecules analyzed: GFP, n=13925; GFP-CEP104, n=92. Scale bar 2 µm. (C-E) FRAP analysis of CEP104 (C) and EB3 (D) at blocked MT plus ends, or dynamic EB3 at growing plus ends (E). Arrowhead marks point of photobleaching in representative kymographs, scale bars 2 µm and 60 s (C), 2 µm and 10 s (D, E). Plots show averaged curves with exponential fit. Number of FRAP measurements: CEP104 n=17; stationary EB3 n=14; dynamic EB3 n=10. Error bars represent s.e.m..

**Figure 3.**
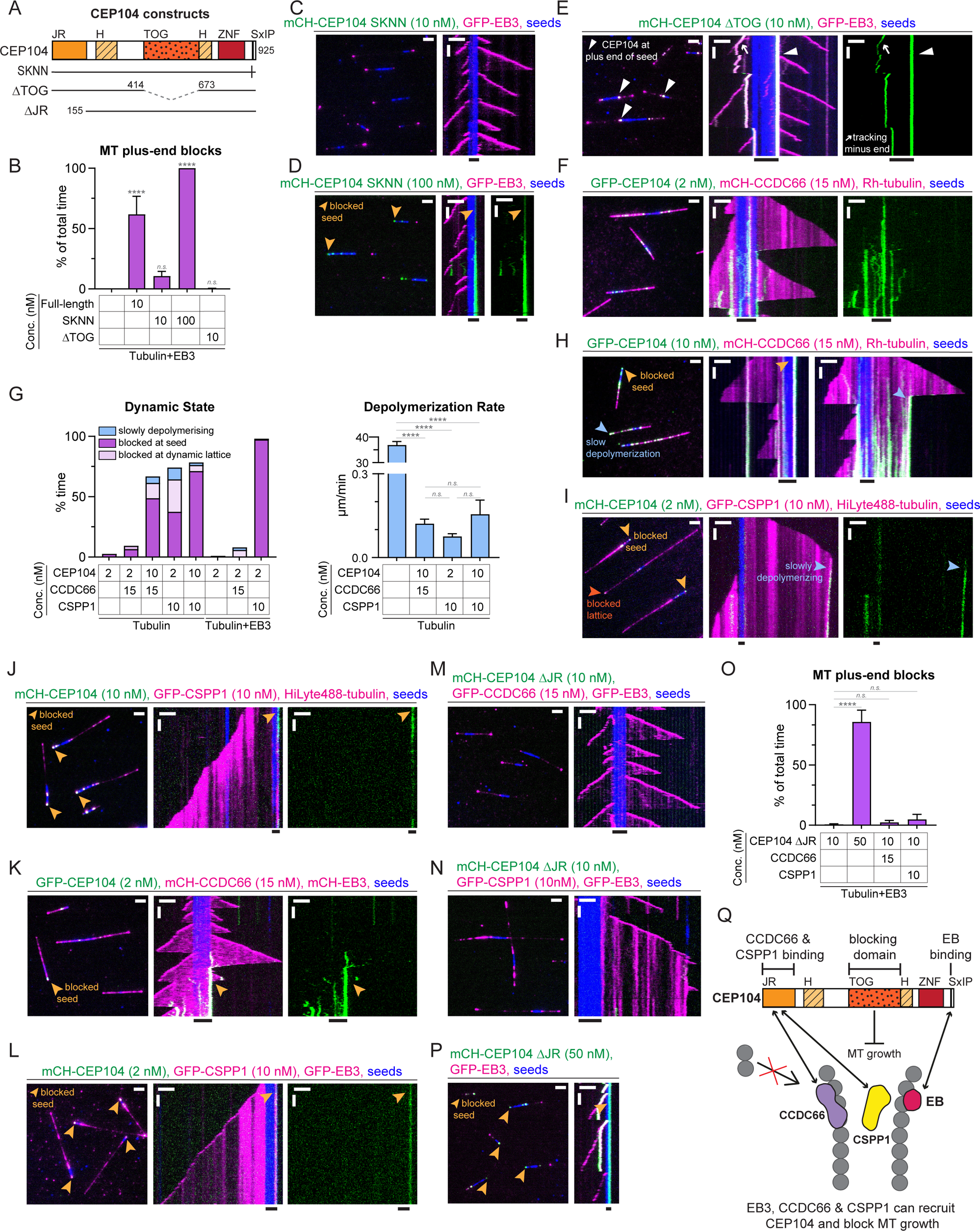
Plus-end blocking by CEP104 is potentiated by EB3, CCDC66 and CSPP1. (A) Schematic representation of different CEP104 constructs; jelly-roll, JR; α-helical domain, H; zinc finger, ZNF. (B) Percentage of time MT plus ends spent blocked in the presence of 20 nM EB3 in combination with CEP104 constructs (from kymographs shown in C-E, 1C, 1L). Bars represent averaged means from three independent experiments. Error bars represent s.e.m.. (C-F) Fields of view (left, scale bar 2 µm) and kymographs (right, scale bars 2 µm and 60 s) illustrating MT dynamics from GMPCPP-stabilized seeds with either 15 µM tubulin supplemented with 3% HiLyte488 or Rhodamine-labelled tubulin or, 20 nM GFP- or mCherry-EB3 and indicated concentrations and colors of ciliary tip module constructs. Orange arrowheads, blocked plus ends; white arrowheads, CEP104 ΔTOG at plus ends of seeds; white arrows, CEP104 tracking the minus end. (G) Parameters of MT plus-end dynamics in the presence of 15 µM tubulin alone or with 20 nM EB3 in combination with indicated concentrations of ciliary tip module proteins (from kymographs shown in F, H-L, S2B). For dynamic state bars represent averaged means from three independent experiments. For depolymerization rate bars represent pooled data from three independent experiments: 2 nM CEP104, n=133; 2 nM CEP104 with 15 nM CCDC66, n=23; 10 nM CEP104 with 15 nM CCDC66, n=122; 2 nM CEP104 with 10 nM CSPP1, n=69; 10 nM CEP104 with 10 nM CSPP1, n=99; EB3 with 2 nM CEP104, n=1; EB3 with 2 nM CEP104 and 15 nM CCDC66, n=63; EB3 with 2 nM CEP104 and 10 nM CSPP1, n=116. (H-N) Fields of view (left, scale bar 2 µm) and kymographs (right, scale bars 2 µm and 60 s) illustrating MT dynamics from GMPCPP-stabilized seeds with either 15 µM tubulin supplemented with 3% HiLyte-488- or Rhodamine-labelled tubulin, or 20 nM GFP or mCherry-EB3 and indicated concentrations and colors of ciliary tip module constructs. Light orange arrowheads, blocked seeds; dark orange arrowhead, blocked lattice; blue arrowheads, slow depolymerization of plus ends. (O) Percentage of time of MT plus ends spent blocked at the seed in the presence of 20 nM EB3 in combination with CEP104 ΔJR and other ciliary tip module proteins (from kymographs shown in M, N, P, S2B). Bars represent averaged means from three independent experiments. (P) Fields of view (left, scale bar 2 µm) and kymographs (right, scale bars 2 µm and 60 s) illustrating MT dynamics from GMPCPP-stabilized seeds with 20 nM GFP- and 50 nM mCherry-CEP104 ΔJR. Orange arrowheads, blocked plus ends. (Q) Schematic model showing CEP104 domain functions and interactions with other ciliary tip module proteins. Jelly-roll, JR; zinc finger, ZNF; α-helical domain, H. For (B, G and O) error bars represent s.e.m. ****, p<0.0001; n.s., not significant; Kruskal-Wallis test followed by Dunńs post-test.

Whereas the SxIP motif was not essential for plus-end blocking by CEP104, the single TOG domain of CEP104 was required to inhibit MT growth. CEP104 lacking the TOG domain (Fig. 3A and S2A) could still be recruited to MTs by EB3, but no plus-end blocking was observed (Fig. 3B and E). In the presence of EB3, CEP104 ΔTOG could bind to growing MT ends and accumulate at the border between the seed and the plus-end-grown lattice (Fig. 3E), a localization for which we currently have no explanation.

Since recent studies described functional and biochemical interactions between CEP104, CCDC66, CSPP1 and ARMC9 (Frikstad et al., 2019; Latour et al., 2020; Odabasi et al., 2023), we next tested *in vitro* the combinations of these proteins. We found that in the presence of 15 nM CCDC66, already 2 nM CEP104 was sufficient to occasionally block MT plus end outgrowth from the seed (Fig. 3F and G). 2 nM CEP104 on its own, or in the presence of EB3, or together with ARMC9, showed hardly any MT binding (Fig. 3G and S2B). However, CEP104 driven growth inhibition at MT seeds became much more common at 10 nM CEP104 with 15 nM CCDC66, and in these conditions, CEP104 and CCDC66 could also strongly inhibit depolymerization of dynamic MTs by either pausing the plus ends or reducing their shrinkage rate from 36.91±1.34 µm/min to 0.12±0.02 µm/min (Fig. 3G and H). Similar effects were observed when either 2 nM or 10 nM CEP104 was combined with 10 nM CSPP1: both growth inhibition at the seed and at the dynamic plus ends, as well as slow depolymerization were observed (Fig. 3G, I and J). These data indicate that CEP104 is not just a MT growth inhibitor or a depolymerase that can prevent tubulin addition to the plus ends, but also a specific MT stabilizing factor that can inhibit tubulin loss from MT plus ends lacking a GTP cap. The addition of EB3 had no strong impact on MT dynamics when combined with 2 nM CEP104 and 15 nM CCDC66 (Fig. 3F, G and K), but potentiated seed blocking when included with 2 nM CEP104 and 10 nM CSPP1, as most MTs were blocked at the seed in these conditions (Fig. 3G, I and L).

Based on our own and previously published co-immunoprecipitation experiments, the interactions of CCDC66 and CSPP1 with CEP104 depended on its jelly-roll (JR) domain (Fig. S2C) (Frikstad et al., 2019; Odabasi et al., 2023). The deletion of this domain abolished the ability of both proteins to recruit CEP104 to MTs (Fig. 3M-O and S2D). However, CEP104 ΔJR could still block seed outgrowth in the presence of EB3 (Fig. 3O and P), though a higher concentration of CEP104 ΔJR compared to the full-length CEP104 was needed to achieve significant plus-end blocking (50 nM versus 10 nM, compare Fig. 1D and L to Fig. 3O, P and S2B), indicating that the JR domain might directly or indirectly contribute to MT blocking.

Taken together, our results demonstrate that CEP104 can stably and specifically associate with MT plus ends and inhibit both tubulin addition and removal from these ends in a TOG domain-dependent manner. However, the affinity of CEP104 for MTs is rather low, and its activity is strongly enhanced by association with its binding partners EB3, CCDC66 and CSPP1 that recruit it to MTs. Since EB3 binds to the outer MT surface (Maurer et al., 2012) and the same might be true for CCDC66 as it can bundle MTs (Batman et al., 2022), whereas CSPP1 associates with the MT lumen (van den Berg et al., 2023), CEP104 might be positioned at protofilament ends to allow access to both surfaces of the MT wall (Fig. 3Q).

### The rescue activity of TOGARAM1 depends on TOG3 and TOG4 domains

Having characterized the activity of CEP104, we next turned to the other TOG domain component of the ciliary tip module, TOGARAM1, which contains two pairs of TOG domains (TOG1-TOG4) connected by a long linker (Fig. 4A). Single molecule counting experiments showed that it is a monomer (Fig. 4B), similar to other MT regulators with multiple TOG domains, XMAP215 (Brouhard et al., 2008) and CLASP2 (Aher et al., 2018). A previous study showed that isolated TOG2 and TOG4, but not TOG1 and TOG3 promoted tubulin polymerization in light scattering assays (Das et al., 2015). To test how these domains contribute to the rescue activity of the full-length protein, we took advantage of previously generated TOGARAM1 mutants where the tubulin-binding surfaces of the individual TOG domains were disrupted by point substitutions within the intra-HEAT loop of HEAT repeat A (Das et al., 2015) (Fig. 4A). Although there is no direct interaction between TOGARAM1 and EB3, we included EB3 in our *in vitro* reconstitution assays as EB3 increases catastrophe frequency and therefore facilitates observing rescues (Fig 1F) (Komarova et al., 2009). We found that simultaneous mutation of TOG3 and TOG4 (TOGARAM1 123’4’), but not TOG1 and TOG2 (TOGARAM1 1’2’34) abolished the rescue activity, although both mutants could still bind to MTs at 10 nM concentration (Fig. 4A, C-E and S3). This was different from previous observations in cells, where the double TOG3-TOG4 mutant (TOGARAM1 123’4’) could not bind to MTs (Das et al., 2015). TOG3 and TOG4 showed redundancy within the full-length protein because single mutations in each one of them did not abolish rescues (Fig. 4A, C, F, G and S3). TOG1 and TOG2 could be deleted without impairing rescue activity (the construct L-TOG-34, Fig. 4A, C, H and S3), and the combination of these two domains (TOG-12) without the adjacent linker region did not bind to MTs even at 200 nM (Fig. 4A, I and S3). TOG-34 did not visibly associate with MTs at 10 nM, but at 200 nM it bound to MTs and induced rescues (Fig. 4A, C, J, K and S3). Since the construct containing TOG3 and TOG4 domains together with adjacent linker (L-TOG-34) was active in our assays already at 10 nM, in agreement with previous findings, the linker region of TOGARAM1 contributes to MT affinity (Das et al., 2015). All tested constructs mildly reduced MT growth rate and TOG-34 constructs with or without the adjacent linker mildly suppressed catastrophes (Fig. 4C). Our results demonstrate that, on its own, TOGARAM1 has relatively mild effects on MT dynamics, which rely in its two C-terminal TOG domains.

**Figure 4.**
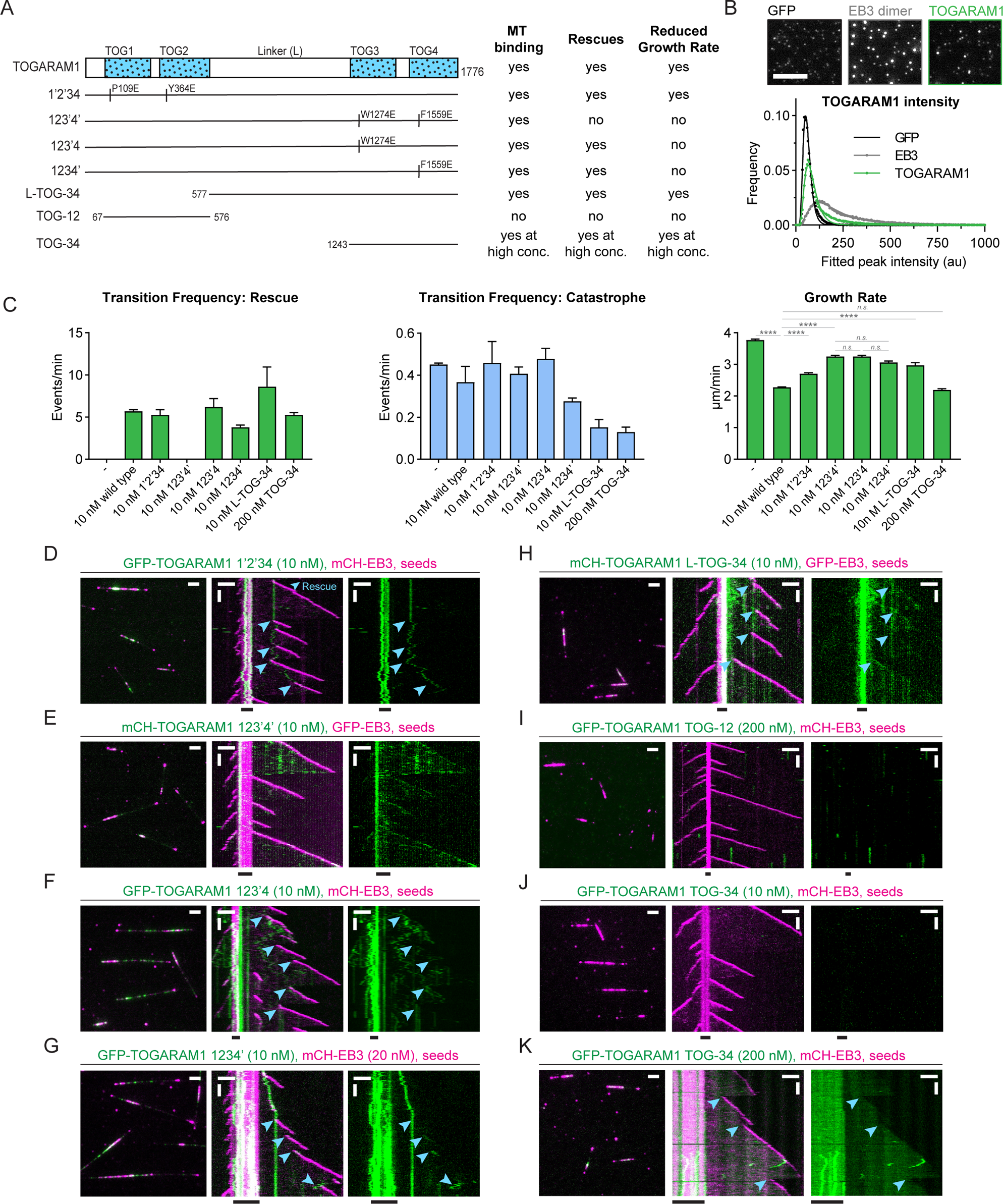
The rescue activity of TOGARAM1 depends on the TOG3 and TOG4 domains. (A) Schematic representation of different TOGARAM1 constructs and summary table highlighting their effects on MT dynamics. Vertical lines indicate point mutations predicted to ablate tubulin-binding activity. (B) Fields of view (top) and histogram plot (bottom) of fluorescent intensities of single GFP molecules, EB3 dimers and TOGARAM1 molecules were immobilized in separate chambers of the same coverslip. Number of molecules analyzed: GFP, n=57865; EB3, n=73074; TOGARAM1, n=74306. Scale bar, 2 µm. (C) Parameters of MT plus-end dynamics in the presence of 20 nM EB3 in combination with indicated concentrations of TOGARAM1 constructs (from kymographs shown in D-H, K, 1C, 1P). For growth rates bars represent pooled data from three independent experiments, total number of growth events: EB3 alone, n=938; EB3 with TOGARAM1, n=861; EB3 with 1’2’34, n=350; EB3 with 123’4’, n=337; EB3 with 123’4, n=560; EB3 with 1234’, n=253; EB3 with L-TOG-34, n=131; EB3 with 200nM TOG-34, n=150. For transition frequencies bars represent average means from three independent experiments. Error bars represent s.e.m., ****, p<0.0001; n.s., not significant; Kruskal-Wallis test followed by Dunńs post-test. (D-K) Fields of view (left, scale bar 2 µm) and kymographs (right, scale bars 2 µm and 60 s) illustrating MT dynamics from GMPCPP-stabilized seeds with 20 nM GFP- or mCherry-EB3 and indicated concentrations and colors of TOGARAM1 constructs. Blue arrowheads, rescues.

### ARMC9 colocalizes with TOGARAM1 and CSPP1 and enhances their effects on MT dynamics

We next set out to characterize the function of ARMC9. Single molecule counting experiments showed that ARMC9 forms dimers dependent on its central helical region (Fig. S4A and B). As shown above, ARMC9 does not bind to MTs on its own nor in the presence of CEP104 and EB3 (Fig. 1G, H and S2B). Consistently, no direct interaction has been described for ARMC9 and CEP104, even though they were previously shown to weakly co-immunoprecipitate with each other (Latour et al., 2020), an observation we were not able confirm (Fig. S4C). We could confirm by co-immunoprecipitation the interaction between ARMC9 and CCDC66, however when this combination was tested in reconstitution experiments we did not see any recruitment of ARMC9 to the MT lattice (Fig. S4C and D). We also confirmed by co-immunoprecipitation the interaction between ARMC9 and TOGARAM1 (Fig. S4C) and refined the previous mapping of this interaction (Latour et al., 2020) by showing that it requires the armadillo (ARM) repeats of ARMC9, but not its helical domain responsible for dimerization, and the tubulin-binding surface of the TOG2 domain of TOGARAM1 (Fig. 4A, S4A, E and F). We also tested the impact of different point mutations identified in Joubert syndrome patients on the interaction of ARMC9 and TOGARAM1 (Latour et al., 2020; Van De Weghe et al., 2017), and found that all tested mutations in the TOG2 domain (all located in the first HEAT repeat) and one of the several known mutations in the ARM domain (G492R) perturb binding (Fig. S4A, G and H), supporting the functional importance of this interaction.

In the *in vitro* assays, TOGARAM1 could recruit ARMC9 to MTs, and the colocalization between the two proteins was particularly prominent in particles that could stationarily bind to or diffuse on MTs (Fig. 5A). As expected from the co-immunoprecipitation experiments, this colocalization was not perturbed by the 123’4’ mutant of TOGARAM1 (Fig. 4A, 5B) but was abolished by the 1’2’34 mutant (Fig. 4A, 5C). The brightness of ARMC9 and TOGARAM1 in MT-bound particles increased over time, indicating binding of additional molecules (Fig. 5A and B). This binding was likely driven by ARMC9, as it was not observed for TOGARAM1 alone (Fig. 1P). Presence of ARMC9 did not drastically alter MT growth rates but did cause a roughly two-fold increase in the rescue activity of TOGARAM1, which was still dependent on TOG3 and TOG4 (Fig. 5A, B, and D). In contrast to ARMC9, the addition of CCDC66 did not alter the effects of TOGARAM1 on MT dynamics, and no colocalization between CCDC66 and TOGARAM1 was observed (Fig. S4D).

**Figure 5.**
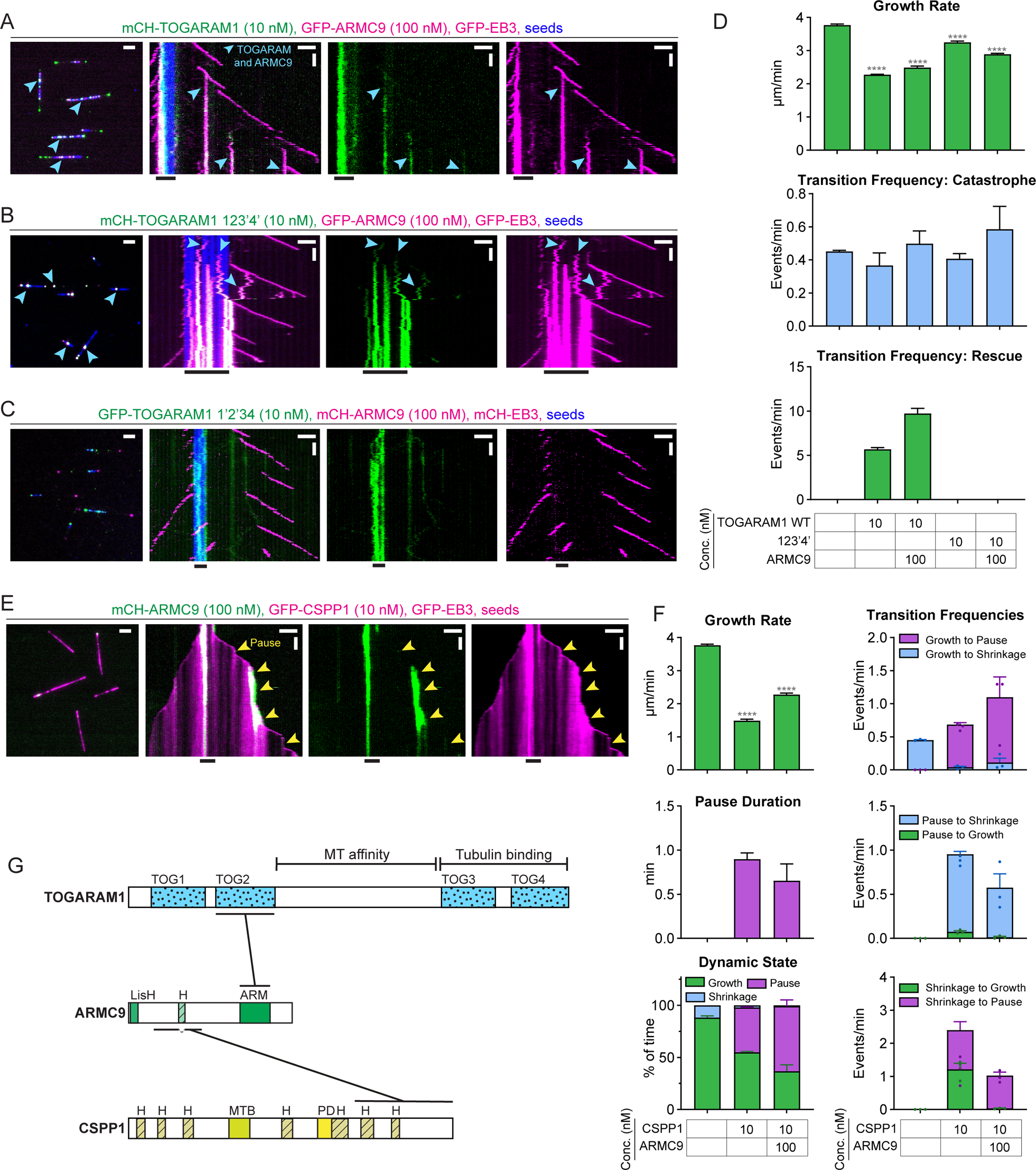
ARMC9 colocalizes with TOGARAM1 and CSPP1 and enhances their effects on MT dynamics. (A-C) Fields of view (left, scale bar 2 µm) and kymographs (right, scale bars 2 µm and 60 s) illustrating MT dynamics from GMPCPP-stabilized seeds with 20 nM GFP- or mCH-EB3 and indicated concentrations and colors of TOGARAM1 constructs and ARMC9. Blue arrowheads, ARMC9 and TOGARAM1 colocalization. (D) Parameters of MT plus-end dynamics in the presence of 20 nM EB3 in combination with indicated concentrations of TOGARAM1 constructs and ARMC9 (from kymographs shown in A-B, 1C, 1P, 4E). For growth rates bars represent pooled data from three independent experiments, total number of growth events: EB3 alone, n=938; EB3 with TOGARAM1, n=861; EB3 with TOGARAM1 and ARMC9, n=342; EB3 with 123’4’, n=337; EB3 with 123’4’ and ARMC9, n=325. For transition frequencies bars represent averaged means from three independent experiments. Error bars represent s.e.m.. ****, p<0.0001; n.s., not significant; Kruskal-Wallis test followed by Dunńs post-test. (E) Fields of view (left, scale bar 2 µm) and kymographs (right, scale bars 2 µm and 60 s) illustrating MT dynamics from GMPCPP-stabilized seeds with 20 nM GFP-EB3 and ciliary tip module proteins indicated. Yellow arrowheads, pauses. (F) Parameters of MT plus-end dynamics with 20 nM EB3 in combination with indicated concentrations of ciliary top module proteins (from kymographs shown in E, 1C, 1N). For growth rate and pause duration bars represent pooled data from three independent experiments, total number of growth events, pauses: EB3 alone, n=938, 0; EB3 with CSPP1, n=1715, 273; EB3 with CSPP1 and ARMC9, n=627, 315. For transition frequencies and dynamic state bar represent averaged means from three independent experiments. Error bars represent s.e.m., ****, p<0.0001; n.s., not significant; Kruskal-Wallis test followed by Dunńs post-test. (G) Schematic model showing TOGARAM1, ARMC9, and CSPP1 interaction domains; MTB; pause domain, PD; α-helical domain, H.

We observed co-immunoprecipitation of ARMC9 and CSPP1 (Fig. S5A and B), confirming previous observations (Latour et al., 2020). The interaction required the C-terminal part of CSPP1 and the linker regions surrounding the central helical domain of ARMC9, though not the helical domain itself (Fig. S5A and B). *In vitro* reconstitution assays showed that CSPP1 could trigger ARMC9 accumulation on MTs in the vicinity of the growing MT ends (Fig. 5E) and mildly increased the percentage of time CSPP1-induced MT pausing (Fig. 5F).

We also tested whether CSPP1 and TOGARAM1 can bind to each other but detected no co-immunoprecipitation (Fig. S5C), and CSPP1 displayed no colocalization with either TOGARAM1 or CCDC66 in the *in vitro* assays (Fig. S5D). Altogether, our results indicate that CCDC66 does not colocalize with either TOGARAM1 or CSPP1 in reconstitution assays, whereas ARMC9 can bind to both of these proteins through two distinct domains and serve as a scaffold mildly enhancing their effects on MT dynamics (Fig. 5G).

### Combined action of ciliary tip proteins drives slow processive MT growth

To complete the exploration of all pairwise combinations of the studied ciliary tip proteins, we examined the joint effects of CEP104 and TOGARAM1. The two proteins coprecipitated with each other in a manner dependent on the zinc finger domain of CEP104 and the linker region of TOGARAM1 (Fig. S6A and B). When they were combined on dynamic MTs in the presence of EB3, there were periods of pausing both at the plus end of the seeds and of the dynamic lattice (Fig 6A-C). However, these pauses were transient and followed by growth events (Fig 6D and E), something we did not see with any other CEP104-ciliary tip protein combination. Furthermore, the two proteins together induced periods of very slow MT polymerization with the rates of 0.12±0.01 µm/min, ∼20 times slower than 2.27±0.02 µm/min observed with TOGARAM1 alone (Fig. 6A, C and D). CEP104 and TOGARAM1 can thus interact through regions that do not engage their tubulin-binding domains and together impose slow MT polymerization (Fig. 6F).

**Figure 6.**
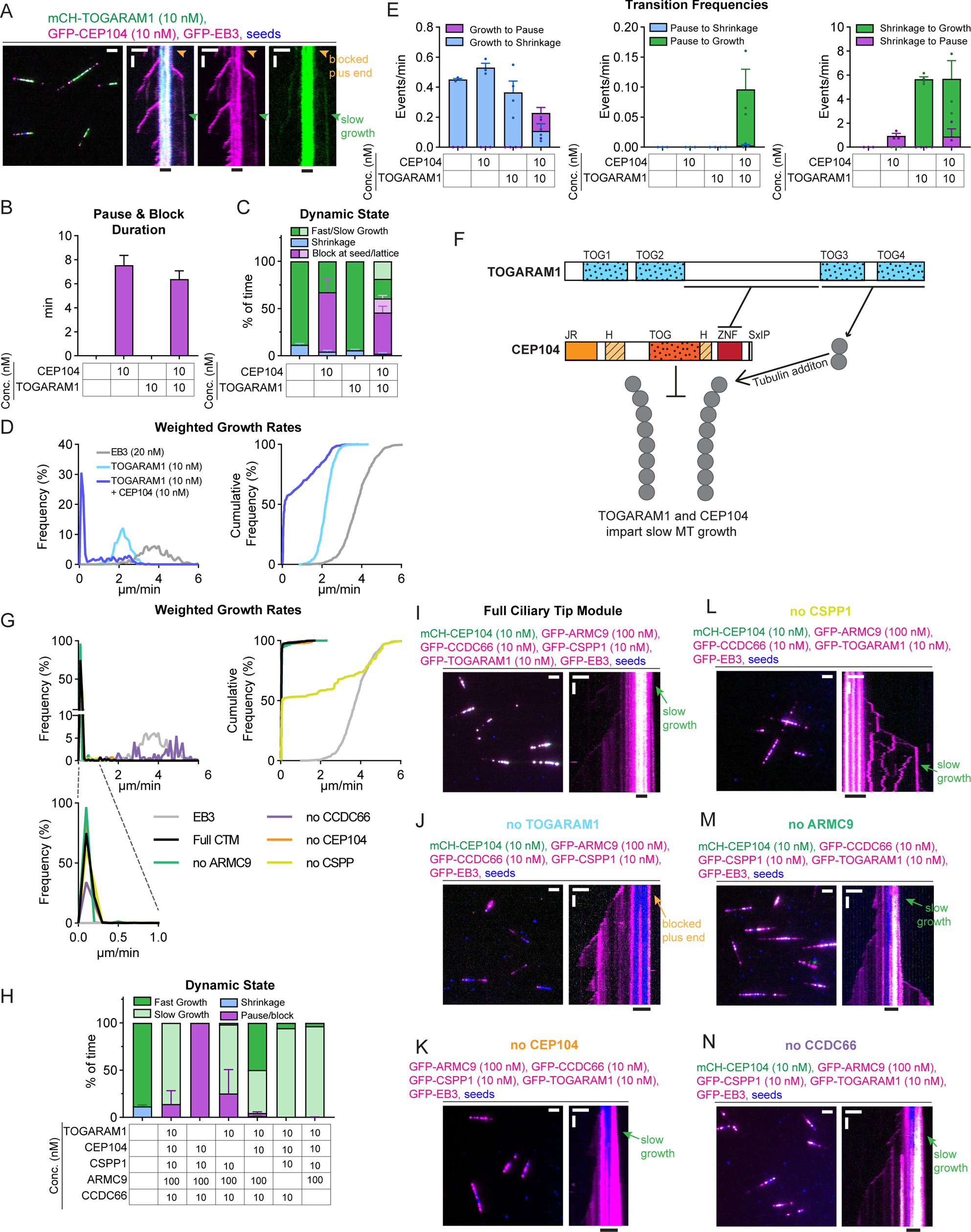
Combined action of ciliary tip proteins drives slow processive MT growth. (A) Fields of view (left, scale bar 2 µm) and kymograph (right, scale bars 2 µm and 60 s) illustrating MT dynamics from GMPCPP-stabilized seed with GFP-EB3, mCherry-TOGARAM1, and GFP-CEP104. Orange arrowheads, blocked plus end; green arrowheads, slow MT growth. (B-E) Parameters of MT plus-end dynamics in the presence of EB3 in combination with indicated concentrations of ciliary tip module proteins (from kymographs shown in A, 1C, 1L, 1P). For pause & block duration (B) and growth rates (D) data was pooled from three independent experiments, total number of growth events, pauses: EB3 alone, n=938, 0; EB3 with CEP104, n=213, 101; EB3 with TOGARAM1, n=861, 0; EB3 with CEP104 and TOGARAM1, n=373, 140. Bars represent mean + s.e.m.. For dynamic state (C) and transition frequencies (E) bars represent averaged means from three independent experiments. Error bars represent s.e.m. (F) Schematic model showing interactions between CEP104, TOGARAM1 and tubulin. Jelly-roll, JR; zinc finger, ZNF; α-helical domain, H. (G-H) Weighted growth rates (G) and dynamic state (H) for combinations of ciliary tip module proteins indicated (from kymographs I-N, and 1C). For growth rates (G) data was pooled from three independent experiments, total number of growth events: EB3 alone, n=938; entire ciliary tip module, n=70; no ARMC9, n=81; no CCDC66, n=83; no CEP104, n=41; no CSPP1, n=141; no TOGARAM1, n=0. For dynamic state (H) bars represent or averaged means from two independent experiments, error bars represent s.e.m.. (I-N) Fields of view (left, scale bar 2 µm) and kymographs (right, scale bars 2 µm and 60 s) illustrating MT dynamics from GMPCPP-stabilized seeds with 20 nM GFP- or mCherry-EB3 and indicated concentrations and colors of ciliary tip module proteins indicated. Orange arrow, blocked plus end; green arrows, slow plus-end growth.

Next, we combined all five ciliary tip module components in the presence of EB3. Strikingly, in these conditions, all MTs displayed slow and highly processive plus-end polymerization with occasional pausing, with an average rate of 0.19±0.04 µm/min, while minus ends could still undergo phases of growth and shrinkage (Fig. 6G-I and S6C). To test which proteins are essential for this dynamic state, we repeated the assays while systematically leaving out each of the ciliary tip proteins. Omitting TOGARAM1 abolished slow growth completely, as all MT plus ends were paused (Fig. 6H and J). This was not surprising, because MT behavior in these conditions was dominated by the two pausing factors, CEP104 and CSPP1. This indicates that an important function of TOGARAM1 is to overcome MT growth inhibition imposed by CEP104 or CSPP1. Leaving out CEP104 had no major effect, except for a slight increase in pausing and very occasional periods of fast growth (Fig. 6G, H and K), whereas fast elongation became much more common in the absence of CSPP1 (Fig. 6G, H, and L). Omitting either ARMC9 or CCDC66 had no major effect except that no pausing was observed, and occasional episodes of fast growth were detected (Fig. 6G, H, M and N). No MT catastrophes or shrinkage were observed in any of these conditions (Fig. 6H). Altogether, ciliary tip module components stabilize MTs by preventing their depolymerization and promoting robust but very slow tubulin addition.

### Ciliary tip proteins form cork-like densities at MT plus ends and reduce protofilament flaring

To better understand the localization of the ciliary tip module on MT ends, we turned to cryo-electron tomography (cryo-ET). We reconstructed 3D volumes containing MTs grown with either EB3 alone or in the presence of the entire ciliary tip module. To best match our *in vitro* reconstitution conditions, samples were incubated at 30 °C with GMPCPP-stabilized seeds and 15 µM tubulin for 10 minutes before being added to holey carbon-coated copper grids and vitrified. Based on our analysis of MT dynamics we presumed that they were mostly fast growing for EB3 controls and elongating slowly with a few paused ends for ciliary tip module samples (Fig. 6H). MTs grown in the presence of the ciliary tip module showed densities at their ends reminiscent of champagne corks, whereas MTs grown in the presence of only EB3 had no clear density at their ends (Fig. 7A-D, S7A and B). Despite the density for the ciliary tip module appearing quite varied in structure, it was always situated in both the lumen and on the tips of the MT protofilaments (Fig. 7B, D and S7B). We used 2D rotational averaging to resolve MT polarity based on chirality of the cross-section (Bouchet-Marquis et al., 2007), through which we determined that cork-like electron densities were specific for MT plus ends (Fig. 7C, D and S7B). 36 out of 40 captured “corks” where confidently assigned to plus ends, the polarity of the other 4 MTs remains undetermined due to poor tilt-alignment of the tomograms. We performed semi-automated segmentation of the tomographic volumes, both for visualization and segregation of the MT protofilaments and ciliary tip module (Fig. 7E and F). It proved difficult to segregate all individual protofilaments from each other for further analysis due to the limited tilt range. However, the majority of protofilaments were individually resolved, and separation between the ciliary tip module and protofilaments was often clear. Extensive contacts on the luminal surface of tubulin and the exposed protofilament ends were observed, while the exterior surface of the MT remained largely unbound.

**Figure 7.**
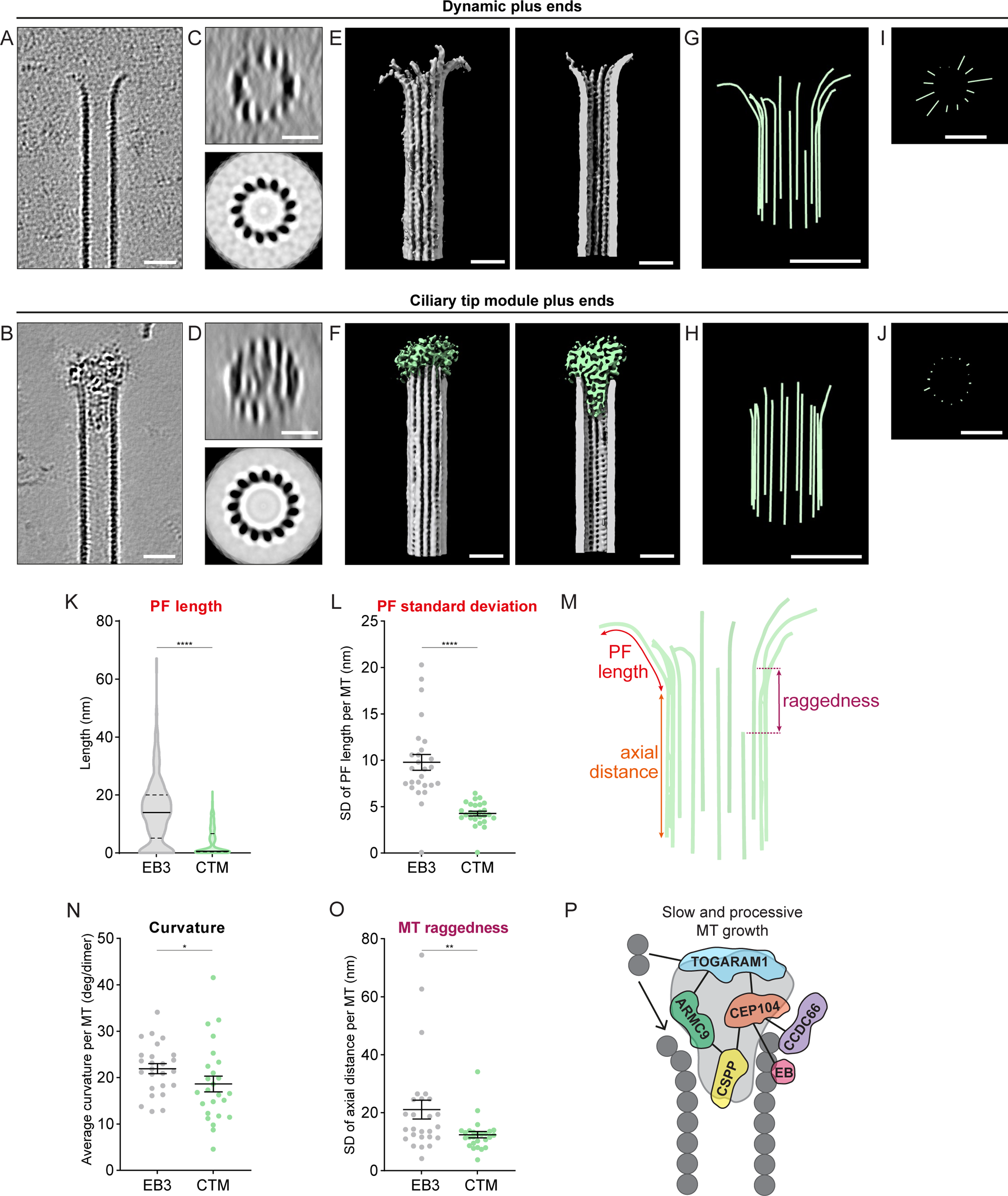
Ciliary tip proteins form cork-like densities at MT plus ends and reduce protofilament flaring. (A, B) Slices (4.3 nm thick) of denoised tomograms showing MT plus ends grown in presence of EB3 alone (A) or ciliary tip module and EB3 (B). Scale bars 25 nm. (C, D) 8nm thick transverse cross-section of the corresponding MTs in panels A and B. The transverse cross-section (top) is accompanied by rotational average (bottom) to indicate MT polarity. Scale bars 25 nm. (E, F) 3D rendered segmentation volume of a free/dynamic MT plus end and ciliary tip module-plugged plus end. The MT is shown in grey and the ciliary tip module density in green. (G-J) 3D model of manually traced protofilament shapes (G, H), accompanied by their transverse cross-sections (I, J). (K) Distribution of protofilament length measured from the last segment within the MT wall until the tip of the protofilament. Total number of protofilaments analyzed; EB3, n=329; CTM, n=352. Lines represent median (solid) and quartiles (dotted). ****, p<0.0001; Mann-Whitney test. (L) Protofilament standard deviation per MT. Total number of MTs analyzed: EB3, n=26; ciliary tip module, n=26. Bars represent mean, with error bars representing s.e.m., ****, p<0.0001 (Mann-Whitney test). (M) Schematic representation of parameters that were obtained from manual tracing of MT plus-end protofilaments. (N) Average local protofilament curvature per MT. Total number of MTs analyzed: EB3, n=25; ciliary tip module, n=25. Bars represent mean, with error bars representing s.e.m., *, p<0.05; Mann-Whitney test. (O) MT raggedness plotted as standard deviation of protofilament axial distance of last segment within the wall per MT. Total number of MTs analyzed: EB3, n=26; ciliary tip module, n=26. Bars represent mean, with error bars representing s.e.m., **, p<0.01; Mann-Whitney test. (P) Schematic model of the ciliary tip module at a MT plus end.

We turned to manual tracing to analyze how the ciliary tip module may be altering MT tip architecture, and how this architecture deviates from more typical MT architecture associated with fast dynamics (Fig. 7G-J, S7A and B). We first looked at how flared the MT protofilaments were by measuring both protofilament length (length of curved segment) and protofilament standard deviation per MT (standard deviation of the distance between first point that deviated from being completely straight to the end of the protofilament, see materials and methods) (Fig. 7K-M). As expected, MT plus ends grown with only EB3 showed primarily flared protofilaments (Gudimchuk et al., 2020; McIntosh et al., 2018) (Fig. 7A, K, L and S7A), whereas plus ends “corked” with ciliary tip module proteins had a dramatic decrease in both protofilament length and standard deviation in protofilament length per MT (Fig. 7B, K, L and S7B), as well as a mild decrease in average local curvature (Fig. 7N). Finally, MT raggedness, defined as the standard deviation in the axial distance along the MT of the first point for each protofilament that deviated from being completely straight (Fig. 7M), was modestly but significantly decreased in MT plus ends “corked” by ciliary tip module proteins (Fig. 7O). Together our results indicate a potential mechanism where the ciliary tip module acts from both the lumen and tips of MTs to stabilize slow and processive elongation (Fig 7P).

## Discussion

Ciliary tip proteins play important roles in controlling the structure and therefore function of motile and primary cilia. Mutations in these proteins cause various ciliopathies often resulting from shorter cilia causing defects in cilia-dependent signaling pathways (Frikstad et al., 2019; Latour et al., 2020; Louka et al., 2018; Odabasi et al., 2023; Perlaza et al., 2022). Axonemal MTs are stable and elongate very slowly without undergoing long depolymerization events. Here, we have reconstituted these behaviors *in vitro* with five components of the ciliary tip module. We found that collectively, these proteins robustly impose very slow and processive MT growth. Furthermore, elongation rates measured in our assays, 0.19±0.04 µm/min, fall within the range of those measured for the initial elongation of regenerating flagella in single-celled organisms such as *Chlamydomonas reinhardtii* (0.08-0.40 µm/min) (Marshall et al., 2005; Rosenbaum and Child, 1967; Rosenbaum et al., 1969; Witman, 1975), though these rates are reduced when flagella reach their normal length. However, it remains to be determined if our measured growth rates are within the range of elongating primary cilia.

Slow growth is an unusual state for dynamic MTs, because it is incompatible with the formation of a long stabilizing GTP cap, which is needed to avoid MT disassembly (Desai and Mitchison, 1997) and promote growth (Wieczorek et al., 2015). Factors controlling slow MT polymerization must thus inhibit tubulin detachment and promote tubulin addition at MT plus ends that have only a very short or no GTP cap. We found that several ciliary tip proteins can perform these functions. Two of them, TOGARAM1 and CEP104, were predicted to be MT polymerases because their individual TOG domains can bind tubulin and enhance tubulin polymerization *in vitro*

(Al-Jassar et al., 2017; Das et al., 2015; Rezabkova et al., 2016; Yamazoe et al., 2020). However, full-length TOGARAM1 and CEP104 do not accelerate tubulin polymerization at freely growing MT plus ends. On the contrary, TOGARAM1 somewhat slows down MT growth rate, and CEP104 potently blocks MT elongation. Still, these proteins are MT stabilizers rather than depolymerases as both can inhibit tubulin detachment from GDP-bound MT ends: TOGARAM1 induces rescues, whilst CEP104 dramatically slows down disassembly of MT plus ends. Moreover, TOGARAM1 can promote tubulin addition and thus serve as a polymerase at paused MT ends, and together CEP104 and TOGARAM1 can induce phases of very slow polymerization.

The processive slow MT growth reconstituted by either CEP104 and TOGARAM1 together, or the ciliary tip module as a whole, is unique to the plus ends of MTs. This is likely driven by the two TOG domain-containing module members, because specific orientation of the TOG domains can position tubulin dimers favorably for binding to protofilament tips at the plus end (Ayaz et al., 2012). Generally, the interaction of TOG domain proteins with MTs depends on positively charged intrinsically disordered regions (Aher et al., 2018; Byrnes and Slep, 2017; Widlund et al., 2011) and additional adaptors, such as EB1 and SLAIN2 (Li et al., 2011; van der Vaart et al., 2011). This is also true for the ciliary TOG domain regulators. TOGARAM1, similar to XMAP215 (Widlund et al., 2011), contains a linker region which promotes binding to MTs *in vitro*, and is further recruited to MTs by ARMC9. CEP104 depends on interactions with several binding partners, EB3, CSPP1 and CCDC66. EB3 is known to bind to the outer MT surface (Maurer et al., 2012), whereas CSPP1 is a luminal protein (van den Berg et al., 2023). The binding mode of CCDC66 has not yet been clarified, but it is likely that some of its parts are present on MT surface as it can bundle MTs *in vitro* (Batman et al., 2022). With binding partners on both sides of the MT wall, CEP104 is in a good position to span the MT tip. TOGARAM1 is likely positioned very close to protofilament ends, since it can catalyze tubulin addition. Therefore, ARMC9 is another good candidate for connecting across MT surfaces as it not only interacts and promotes TOGARAM1 activity but also the pausing behavior of luminal protein CSPP1. Collectively, this fits with the cryo-ET data, which show that the complex of ciliary tip proteins localizes only at the plus ends of MTs and is associated with the luminal side (likely CSPP1) and the tip of MTs (likely CEP104, TOGARAM1 and CCDC66), resembling a cork on a champagne bottle.

The interactions between ciliary tip proteins depend at least partially on non-overlapping domains: for example, CEP104 binds to tubulin through its TOG domain, to EB3 through the SxIP motif within its flexible C-terminal region, to TOGARAM1 through the zinc finger domain and to CCDC66 and CSPP1 through the jelly-roll domain (Fig 3Q, 6F). Similarly, TOGARAM1 interacts with tubulin through TOG3 and TOG4, with ARMC9 through TOG2 and with MTs and CEP104 through its linker region (Fig. 5G). Additionally, Joubert syndrome-linked mutations in TOGARAM1 TOG2 and ARMC9 ARM domains disrupt their interaction, highlighting the functional importance for understanding how the ciliary tip module forms. For several reasons, we do not think the five ciliary tip proteins form a stoichiometric complex but rather a flexible interaction network, the function of which is likely to enhance the accumulation of these proteins at the distal ends of cilia. Firstly, a flexible interaction network could explain why we saw variation in size of “corks” in our cryo-ET data. Additionally, *in vitro*, we saw that ciliary tip module members act in a partially redundant fashion and therefore we cannot be sure that all ciliary tip proteins are present in every “cork”. Leaving individual ciliary tip module proteins out of reconstitutions, whilst making slow growth less robust, had no major effect on MT dynamics. The one exception was TOGARAM1, which was essential for MT elongation; and that aligned well with the observation that overexpression of TOGARAM1 induces longer cilia (Latour et al., 2020). Furthermore, in vertebrate cells, the loss of any of the ciliary tip module proteins results in the generally mild phenotype of shorter cilia, whereas simultaneous loss of several module components has more severe consequences (Das et al., 2015; Frikstad et al., 2019; Latour et al., 2020; Odabasi et al., 2023; Yamazoe et al., 2020). Finally, a loose interaction mode is also in line with the fact that in other species, such as ciliates, these proteins bind to different sites (e.g., A- and B-tubules) and can have opposing functions (Louka et al., 2018).

It appears counterintuitive that deletions or mutations in all vertebrate ciliary tip module proteins cause shorter cilia even though CEP104 and CSPP1 inhibit MT elongation (Das et al., 2015; Frikstad et al., 2019; Latour et al., 2020; Odabasi et al., 2023; Yamazoe et al., 2020). This could be explained by the interplay with additional factors not included in our assays. For example, CEP104 and CSPP1 might counteract MT depolymerase factors, such as kinesin-13, kinesin-18 KIF19A or kinesin-4 KIF7, which are known to act at ciliary tips (Blaineau et al., 2007; He et al., 2014; Niwa et al., 2012). Alternatively, similar to our *in vitro* assays with TOGARAM1, CEP104 and CSPP1 might also cooperate with MT polymerization-promoting factors, such as XMAP215 or CLASP. This idea is supported by CEP104 promoting centriole elongation in flies (Ryniawec et al., 2023), where it is established that Orbit/CLASP and kinesin-13 KLP10A act as positive and negative regulators of fly centriole length, respectively (Shoda et al., 2021).

An important question is the nature of structural changes at MT plus ends associated with slow polymerization. Cryo-ET analysis showed that rapidly growing MTs terminate with strongly flared protofilaments (Gudimchuk et al., 2020; McIntosh et al., 2018), whereas cork-like structures formed by ciliary tip proteins reduce protofilament flaring and at least partially cap the plus ends. Still, in these conditions, MT tips retain conformational variability, with some protofilaments more flared and some more blunt. It is possible that at each given time, the protofilaments that are blunt are occluded by ciliary tip proteins and therefore paused. Despite lacking a GTP cap, these protofilaments don’t depolymerize because ciliary tip regulators keep them together and stabilize them from the luminal side. In this model, the more curved protofilaments are the ones undergoing tubulin addition. This would be consistent with the idea that TOG domains increase local tubulin concentration and deliver ‘curved’ tubulins to protofilament tips (Ayaz et al., 2014). Alternatively, ciliary tip proteins could be responsible for keeping tubulin dimers in flared protofilaments in a bent state, while new dimers are added onto the straighter protofilaments, which slowly forces the corks out of the plus ends as they elongate.

Formation of MT plus-end-specific “corks” that impose blunt protofilament conformation is strikingly similar to the “plugs” observed in reconstitution experiments with the centriolar cap protein CP110 (Ogunmolu et al., 2021). MT stabilization from the luminal side combined with the reduction of protofilament curvature and occlusion of the longitudinal interface of β-tubulin might thus be a common mechanism for generation of very stable and slowly growing MTs. The relevance of such structures is supported by the observations of plugs at the tips of ciliary MTs in cells (Legal et al., 2023; Leung et al., 2021). A mechanism involving the luminal MT surface and occlusion of the longitudinal interfaces of β-tubulin is different from the regulation of much more dynamic cytoplasmic MT ends, which involves factors located on the outer MT surface (Akhmanova and Steinmetz, 2015). Strikingly, the fundamental principles are the same: in both cases, plus-end regulation depends on the balance of growth-promoting and inhibiting activities that are brought together by a multivalent network of interacting proteins.

## Materials and Methods

### DNA constructs, cell lines and cell culture

All proteins were cloned into modified pEGFP-C1 or pmCherry-C1 vectors with a StrepII tag and expressed in HEK293T cells (ATCC). For all module members except TOGARAM1 the human sequence was used, due to technical difficulties the mouse sequence of TOGARAM1 was used, it is 84% identical to human TOGARAM1. HEK293T cells were cultured in DMEM:F10 (1:1) for co-immunoprecipitation assays, or DMEM (Lonza) for protein purification, both were supplemented with 10% fetal calf serum (GE Healthcare Life Sciences) and 1% (v/v) penicillin/streptomycin. All cells were routinely checked for mycoplasma contamination using the MycoAlertTM Mycoplasma Detection Kit (Lonza).

### Co-immunoprecipitation assays

HEK293T cells were transiently transfected with a mix consisting of polyethyleneimine (Polysciences) and constructs for the ciliary tip module proteins. Co-IP was performed with ChromoTek GFP-Trap® Magnetic particles M-270 (Proteintech) following manufacturer’s instructions with company advised buffers. 24 hours post-transfection, cells were harvested in 1x PBS and lysed for 30 minutes on ice in 200 μL lysis buffer (10 mM Tris HCl pH 7.5, 150 mM NaCl, 0.5 mM EDTA, 0.5% IGEPAL CA-630) supplemented with EDTA free protease inhibitor cocktail (Roche) and PhosSTOP™ phosphatase inhibitor cocktail (Roche). Lysate was cleared by 20 min centrifugation at 4 °C at 17,000 x g and diluted with 300 μL dilution buffer (10 mM Tris HCl pH 7.5, 150 mM NaCl, 0.5 mM EDTA) supplemented with EDTA free protease inhibitor cocktail(Roche) and PhosSTOP™ phosphatase inhibitor cocktail (Roche) per sample. Diluted lysate was incubated rotating end-over-end for 45 minutes with wash buffer equilibrated beads. Beads were washed four times with 500 μL wash buffer (10 mM Tris HCl pH 7.5, 150 mM NaCl, 0.5 mM EDTA, 0.05% IGEPAL CA-630) and protein was eluted in 80 μL 2x Laemmli sample buffer (0.125M tris HCl, 20% glycerol, 10% 2-mercaptoethanol, 0.02% Bromophenol blue, 0.2M DTT) and heated at 95 °C for 5 minutes. All steps prior to elution were performed at 4 °C and all reagents and samples were kept on ice.

Prepared samples were run on 8% SDS-page gel and blotted onto Amersham Protran Premium 0.45 μm NC nitrocellulose membrane (Cytiva) by wet transfer at 37 constant volts overnight at 4 °C in transfer buffer (0.2M tris-HCl, 2M glycine, 10% MeOH and 0.01% SDS), or by semi-dry transfer for 1 hour at 0.15 constant Ampère in transfer buffer (0.2M tris-HCl, 2M glycine, 10% MeOH). Membranes were blocked in blocking buffer (2% BSA diluted in 1x PBS 0.05% Tween-20) for 1 hour and sequentially incubated with primary antibodies and secondary antibodies diluted in blocking buffer for 1 hour rotating at room temperature (primary and secondary antibodies) or overnight at 4 °C (primary antibodies only). Blots were thoroughly washed in between and after incubation with antibodies with wash buffer (1x PBS 0.05% Tween-20) and imaged with Odyssey® CLx Infrared Imaging System (LI-COR biosciences) at various exposure times. The following antibodies were used for western blotting: rabbit anti-RFP (1:2000, Rockland immunochemicals), mouse anti-GFP (1:2000, Sigma-Aldrich), donkey anti-mouse IgG (H + L) IRDye® 800CW (1:10000, LI-COR bioscience) and donkey anti-rabbit IgG (H + L) IRDye® 680CW (1:10000, LI-COR bioscience).

### Protein purification from HEK293T cells for in vitro reconstitution assays

HEK293T cells were transiently transfected with a mix consisting of polyethyleneimine (Polysciences) and constructs for one of the ciliary tip module proteins. Protein purification was performed as described before (van den Berg et al., 2023). The cells were harvested 28 hours after transfection in lysis buffer (50 mM HEPES, 300 mM NaCl, 1 mM MgCl_2_, 1 mM DTT, 0.5% Triton X-100, pH 7.4) supplemented with protease inhibitors (Roche) and kept on ice for 15 minutes. After clearance of the debris by centrifugation, the supernatant was incubated with 20 µl StrepTactin beads (GE Healthcare) for 45 min. After several washing steps, five times with a 300 mM salt wash buffer (50 mM HEPES, 300 mM NaCl, 1 mM MgCl_2_, 1 mM EGTA, 1 mM DTT, 0.05% Triton X-100, pH 7.4) and three times with a 150 mM salt wash buffer (50 mM HEPES, 150 mM NaCl, 1 mM MgCl_2_, 1 mM EGTA, 1 mM DTT, 0.05% Triton X-100, pH 7.4), the protein was eluted in elution buffer (50 mM HEPES, 150 mM NaCl, 1 mM MgCl_2_, 1 mM EGTA, 1 mM DTT, 0.05% Triton X-100, pH 7.4, 2.5 mM d-Desthiobiotin (Sigma-Aldrich)). Purified proteins were snap-frozen and stored at −80°C.

### Mass spectrometry

To confirm each protein was purified, and the eluted protein did not contain any contaminants or known interactors that could affect MT dynamics, we performed mass spectrometry (MS) analysis. MS measurements were performed as described previously (van den Berg et al., 2023). The purified protein sample was digested using S-TRAP microfilters (ProtiFi) according to the manufacturer’s protocol. Digested peptides were eluted and dried in a vacuum centrifuge before liquid chromatography-mass spectrometry (LC-MS) analysis. The samples were analyzed by reversed-phase nLC-MS/MS using an Ultimate 3000 UHPLC coupled to an Orbitrap Exploris 480 mass spectrometer (Thermo Scientific). Digested peptides were separated using a 50 cm reversed-phase column packed in-house (Agilent Poroshell EC-C18, 2.7 µm, 50cm x 75 µm) and eluted from the column at a flow rate of 300 nL/min. MS data were acquired using a data-dependent acquisition (DDA) method with a MS1 resolution of 60,000 and mass range of 375-1600m/z. Fragmentation spectra were collected at a resolution of 15,000 using an HCD of 28, isolation window of 1.4m/z, and a fixed first mass of 120m/z. MS/MS fragment spectra were searched using Sequest HT in Proteome Discoverer (Thermo Scientific) against a human database (UniProt, year 2020) that was modified to contain the tagged protein sequence from each ciliary tip module protein and a common contaminants database. Tryptic miss cleavage tolerance was set to 2, fixed modifications were set to cysteine carbamidomethylation, and variable modifications included oxidized methionine and protein N-terminal acetylation. Peptides that matched the common contaminate database were filtered out from the results table.

### In vitro reconstitution assays

#### MT seed preparation

Double-cycled GMPCPP-stabilized MT seeds were used as templates for MT nucleation *in vitro* assays. GMPCPP-stabilized MT seeds were prepared as described before (van den Berg et al., 2023). A tubulin mix consisting of 70% unlabeled porcine brain tubulin, 18% biotin-labeled porcine tubulin and 12% HiLyte-488-, rhodamine-, or HiLyte647-labeled porcine tubulin (all from Cytoskeleton) was incubated with 1 mM GMPCPP (Jena Biosciences) at 37°C for 30 minutes. Polymerized MTs were then pelleted using an Airfuge for 5 min at 199,000 x g and subsequently depolymerized on ice for 20 min. Next, 1 mM GMPCPP was added and MTs were let to polymerize again at 37°C for 30 minutes. Polymerized MTs were again pelleted (as above) and diluted tenfold in MRB80 buffer containing 10% glycerol prior to snap-freezing to store them at −80°C.

#### In vitro reconstitution assays

*In vitro* assays with dynamic MTs were performed as described before (van den Berg et al., 2023). Microscopic slides were prepared by adding two strips of double-sided tape to mount plasma-cleaned glass coverslips. The coverslips were functionalized by sequential incubation with 0.2 mg/mL PLL-PEG-biotin (Susos AG, Switzerland) and 1 mg/mL neutravidin (Invitrogen) in MRB80 buffer (80 mM piperazine-N, N[prime]-bis (2-ethane sulfonic acid), pH 6.8, supplemented with 4 mM MgCl_2_, and 1 mM EGTA). Then, GMPCPP-stabilized MT seeds were attached to the coverslips through biotin-neutravidin interactions. The coverslip was blocked with 1 mg/mL κ-casein before the reaction mix was flushed in. The reaction mix consisted of different concentrations and combinations of fluorescently labeled purified proteins in MRB80 buffer supplemented with 15 µM porcine brain tubulin (100% dark porcine brain tubulin when 20 nM GFP-EB3 or mCherry-EB3 was added, or 97% dark porcine brain tubulin with 3% rhodamine- or HiLyte-488-labeled porcine tubulin), 0.1% methylcellulose, 1 mM GTP, 50 mM KCl, 0.2 mg/mL κ-casein, and oxygen scavenger mix [50 mM glucose, 400 µg/mL glucose oxidase, 200 µg/mL catalase and 4 mM DTT]. This mix was spun down in an Airfuge for 5 min at 119,000 x g before addition to the flow chamber and the flow chamber was sealed with vacuum grease. Microtubules were imaged immediately at 30°C using a total internal reflection fluorescence (TIRF) microscope.

### In vitro assays for cryo-ET

Samples were prepared as above for reconstitution assays with slight modifications. Instead of flow chambers, all steps occur in a tube. Reaction mixes consisted of either just 20nM EB3 or all CTM components (20nM TOGARAM1, 20nM CEP104, 20nM CSPP1, 20nM CCDC66, 200nM ARMC9) with 20nM EB3 in MRB80 buffer supplemented with 15 µM porcine brain tubulin, 1 mM GTP, 0.2 mg/mL κ-casein, and 15 µM DTT. After centrifugation of the reaction mix for dynamic MTs, GMPCPP-stabilized seeds and ProtA-Au^5^ fiducials (1:20) were added, and MTs were left to polymerize for 10-20 min at 30°C in an Eppendorf tube. Subsequently, the sample was gently resuspended and 4 µl was transferred to a glow-discharged holey carbon R2/1 copper grid (Quantifoil Micro Tools) suspended in the chamber of a Vitrobot (Thermo Fisher Scientific). The sample was incubated for 2 min inside the Vitrobot chamber, equilibrated at 30 °C and 95% relative humidity, to allow for potential MT repolymerization after sample transfer to the grid. Subsequently, grids were manually back-blotted for 3 s and plunge-frozen in liquid ethane. Vitrified grids were clipped into auto-grid cartridges and stored in liquid nitrogen until future use.

### Single-molecule fluorescence intensity analysis

Protein samples of GFP, GFP-EB3 and either GFP-CEP104, GFP-TOGARAM1 or GFP-ARMC9 were diluted in PBS and immobilized in adjacent flow chambers of the sample plasma cleaned glass coverslip as described in (Sharma et al., 2016). Flow chambers were then sealed with vacuum grease and immediately imaged using TIRF microscopy. 10-20 images (per condition) of previously unexposed coverslip areas were acquired with 200 ms exposure time.

### CEP104 molecule counting at blocked MT plus ends

To determine the number of CEP104 molecules at blocked MT plus ends we immobilized GFP in one chamber of the same plasma cleaned glass coverslip and *in vitro* reconstitution assay in the adjacent chamber (as described above). Chambers were sealed and immediately imaged using TIRF microscope. First images on unbleached GFP single molecules were acquired, then using the same illumination conditions, images of unexposed GFP-CEP104 bound MTs were acquired.

### Fluorescence Recovery After Photobleaching

For FRAP experiments, *in vitro* reconstitutions were prepared as described above and imaging was conducted using TIRF microscopy. Either focused 488-laser (for GFP-EB3) or 561-laser (for mCherry-CEP104) was used to bleach specific regions of the MT lattice.

### Microscopy

#### TIRF microscopy

*In vitro* reconstitution assays imaged on a previously described (iLas2) TIRF microscope setup (van den Berg et al., 2023). We used the iLas3 system (Gataca Systems (France)) which is a dual laser illuminator for azimuthal spinning TIRF (or Hilo) illumination. This system was installed on Nikon Ti microscope (with the perfect focus system, Nikon), equipped with 489 nm 150 mW Vortran Stradus 488 laser (Vortran) and 100 mW 561 nm OBIS laser (Coherent), 49002 and 49008 Chroma filter sets. Additionally, a CCD camera CoolSNAP MYO (Teledyne Photometrics) was installed and the set up was controlled with MetaMorph 7.10.2.240 software (Molecular Device). To keep the *in vitro* samples at 30 °C, a stage top incubator model INUBG2E-ZILCS (Tokai Hit) was used. The final resolution using CCD camera was 0.045 μm/pixel. For all assays MT dynamics was measured at 200 ms exposure and 3 second intervals for 10 minutes. For EB3 FRAP experiments continuous imaging was used with 100 ms exposure. For kinesin experiments first a 10 minute movie was acquired at 200 ms exposure and 3 second intervals to measure MT dynamics, subsequently Dm-KHC(1-421)-SNAP-6xHis (gift from Kapitein lab, Addgene plasmid #196976) labelled with Alexa647-SNAP dye (NEB) was continuous imaged using 100 ms exposure.

#### Cryo-ET

Vitrified *in vitro* reconstituted MTs were imaged on a Talos Arctica transmission electron microscope (200 keV) (Thermo Fisher Scientific), equipped with a K2 summit direct electron detector (Gatan) and post-column energy filter aligned to the zero-loss peak with 20 keV slit width. Image acquisition was controlled by Serial-EM (Mastronarde, 2005). MT ends were located on TEM overview images, acquired at 4,100x magnification (33.3 Å/pixel). Tilt series were collected at 63,000x magnification (2.17 Å/pixel) with a dose rate of ∼5 e^−^/pixel/s and a total dosage of 100 e^−^/Å^2^. Tilt images were recorded using a dose symmetric scheme (Hagen et al., 2017) over a tilt range of 54° to −54° or 60° to −60° with a tilt increment of 2°, at a defocus target of −3 μm.

### Image analysis

#### Analysis of MT plus-end dynamics in vitro

Analysis of MT plus-end dynamics was performed as described before (van den Berg et al., 2023). Movies of dynamic MTs were corrected for drift, and kymographs were generated using the plugin KymoResliceWide v.0.4 in ImageJ (https://github.com/ekatrukha/KymoResliceWide). MT tips were traced with lines, and measured lengths and angles were used to calculate the MT dynamics parameters such as growth rate and transition events. All events with growth rates faster than 0.02 µm/min and slower than 0.5 µm/min were categorized as slow growing, events faster than 0.5 µm/min were categorized as fast growing, events with shrinkage rates faster than 0.02 µm/min were categorized as shrinking. The events with slower growth rates or faster shrinkage rates than the before mentioned rates were categorized as pause events. Transition frequencies were calculated by dividing the sum of the transition events per experiment by the total time this event could have occurred. Weighted growth rate histograms were calculated by taking individual growth durations and dividing by total growth time.

#### Analysis of FRAP experiments

Data were normalized to 1 for image acquired immediately before the FRAP event and 0 for the image acquired immediately after the FRAP event using the following equation; Normalized intensity=(intensity _t=x_ – intensity _t=1_)(intensity _t=0_ – intensity _t=1_). For EB3, non-linear fit curves were fitted to the post-bleach intensities and half-life calculated using GraphPad Prism.

#### Analysis of single molecule fluorescence intensities

Single molecule fluorescence spots of proteins immobilized directly on the cover glass were detected and fitted with 2D Gaussain function using custom written ImageJ plugin DoM-Utrect (https://github.com/ekatrukha/DoM_Utrecht). Fitted peak intensities were then used to create intensity histograms. CEP104-blocked plus ends were then manually annotated and fitted with 2D Gaussian function using the same plugin.

#### Cryo-ET 3D volume reconstruction and analysis

Tomogram reconstruction and denoising was performed using an in house Python script, as described previously (Chaillet et al., 2023). The tilt series’ frames were initially aligned and dose-weighted using MotionCor2 (Zheng et al., 2017). Subsequent tilt series alignment and tomographic reconstruction for denoising, visualization and analysis were done through AreTomo 1.3.3. Final reconstructed tomographic volumes were created through weighted back-projection and binned with a factor of two. Cryo-CARE 0.2.2 was used for tomogram denoising (Buchholz et al., 2019). For this, even and odd tomographic reconstructions were generated through splitting of the movie frames for each individual tilt into even and odd summed frames with MotionCor2 (Zheng et al., 2017). Alignment parameters were first calculated on the full tilt series and subsequently applied to the even and odd stacks. The cryo-CARE network was trained on 5 tomograms per dataset, and then applied to the entire set.

For visualization purposes, semi-automated segmentation was performed on denoised tomograms using the EMAN2 2.91 tomoseg module (Chen et al., 2017). For this, separate neural networks were trained for the two features of interest: “MT” and “ciliary tip module”. Each feature was included as ‘negative’ training reference, together with background noise and ice contamination, for the neural network training of the other feature. Final visualization and 3D rendering was performed in ChimeraX.

MT ends, both free and CTM-plugged, were only analyzed for protofilament shape in case they were plus ends. MT polarity was determined through 2D rotational averaging with applied symmetry, using the EMAN2 program *e2proc2d*. Manual 3D tracing of protofilament shapes at MT plus ends was performed, as previously described (McIntosh et al., 2018; McIntosh et al., 2020; Ogunmolu et al., 2021), in the IMOD 4.9.6 program *3dmod* (Kremer et al., 1996). Manual tracing was performed on tomographic subvolumes, generated with the *mtrotlong* program of IMOD (4.11.20). Traces of individual protofilaments were saved as separate contours within one object per MT. To satisfy the three-point requirement of *howflared,* for blunt end protofilaments a third point was added one pixel upstream of the second point that indicated the last protofilament segment in the MT wall. Subsequently, protofilament coordinates were extracted using the *howflared* program of IMOD (4.11.20). MT raggedness and protofilament length parameters, as described in the results section, were calculated by *howflared*. Curvature analysis was performed on the *howflared* output with Matlab scripts available at https://github.com/ngudimchuk/Process-PFs.

### Statistical analysis

Statistical analysis was performed using GraphPad Prism 9. Figure legends contain statistical details of each experiment, including the statistical tests used, the number of measurements and the number of experiments.

### Code availability

The python script for automated tomogram reconstruction and cryo-CARE denoising is available online at https://github.com/SBC-Utrecht/cryocare-from-movies. MATLAB scripts for analysis of protofilament tracings in IMOD are available online at https://github.com/ngudimchuk/Process-PFs.

## Acknowledgements

We thank members of the Akhmanova and Howes labs for insightful discussions. We also thank V. Volkov (Queen Mary, University of London) for advice on cryo-ET sample preparation and V. Volkov and N. Gudimchuk (Lomonosov Moscow State University) for help with cryo-ET analysis. Electron imaging was performed at the Utrecht University (UU) Electron Microscopy Centre. We thank Ingr. C. Schneijdenberg and M. Bergmeijer for cryo-ET microscopy support. This work was supported by the European Research Council Synergy grant 609822 to A.A., The Dutch Organisation for Health Research and Development (ZonMw) grant 09120012110085 to R.R. and A.A., Dutch Research Council (NWO) grant OCENW.XL21.XL21.048 to A.A. and S.C.H., and EMBO Long Term Fellowship AFTL 74-2022 to H.A.J.S.

## Author Contributions

H.A.J.S. and C. M. vdB. performed experiments, analyzed data and wrote the paper, R.A.H. performed and analyzed cryo-electron tomography experiments, and wrote the paper, D.S. performed immunoprecipitation experiments, K.E.S. performed and analyzed mass spectrometry experiments, S.H. supervised and analyzed cryo-electron tomography experiments, R.R has generated essential reagents and acquired funding, A.A. coordinated the project, acquired funding and wrote the paper. All authors reviewed and edited the paper.

## Competing financial interests

The authors declare no competing financial interests.

---

**Figure S1 (Related to Figure 1).**
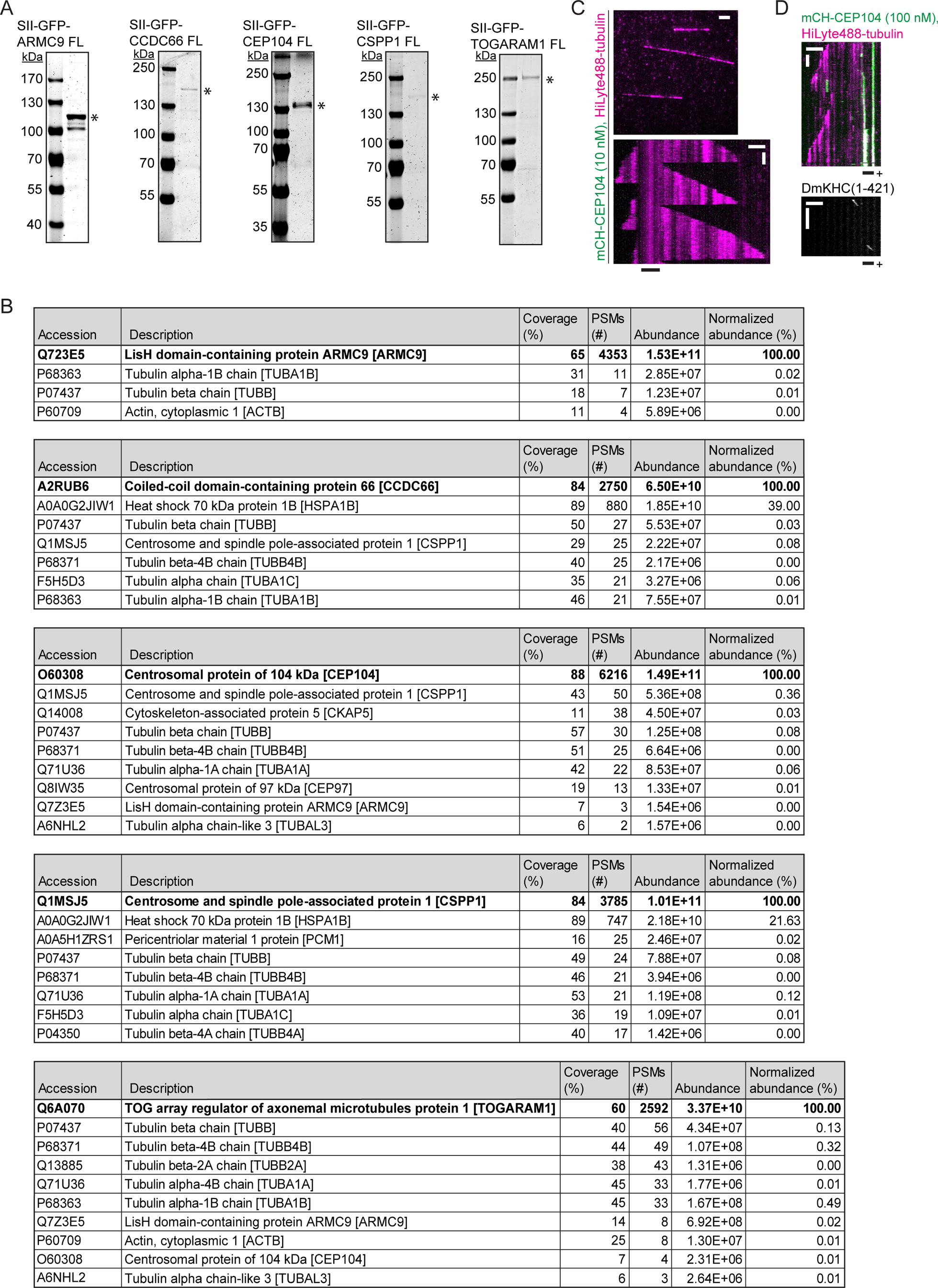
Characterization of ciliary tip module proteins *in vitro*. (A) Analysis of purified GFP-tagged ciliary tip module proteins by SDS-PAGE. Asterisks indicate the full-length protein bands. Protein concentrations were determined from a BSA standard (not shown). (B) Mass spectrometry analysis of purified GFP-tagged ciliary tip module proteins. For all module members except TOGARAM1 the human sequence was used, due to technical difficulties the mouse sequence of TOGARAM1 was used, it is 84% identical to human TOGARAM1. (C) Fields of view (top, scale bar 2 µm) and kymograph (bottom, scale bars 2 µm and 60 s) illustrating MT dynamics from GMPCPP-stabilized seed with HiLyte-488-tubulin and mCherry-CEP104. (D) Kymographs illustrating mobility of DmKHC(1-421) on CEP104-blocked MT labelled with HiLyte-488-tubulin and mCherry-CEP104 (top) and DmKHC (bottom) proving that the blocked end of the MT is the plus end. Scale bars 2 µm and 60 s for both kymographs.

**Figure S2 (Related to Figure 3).**
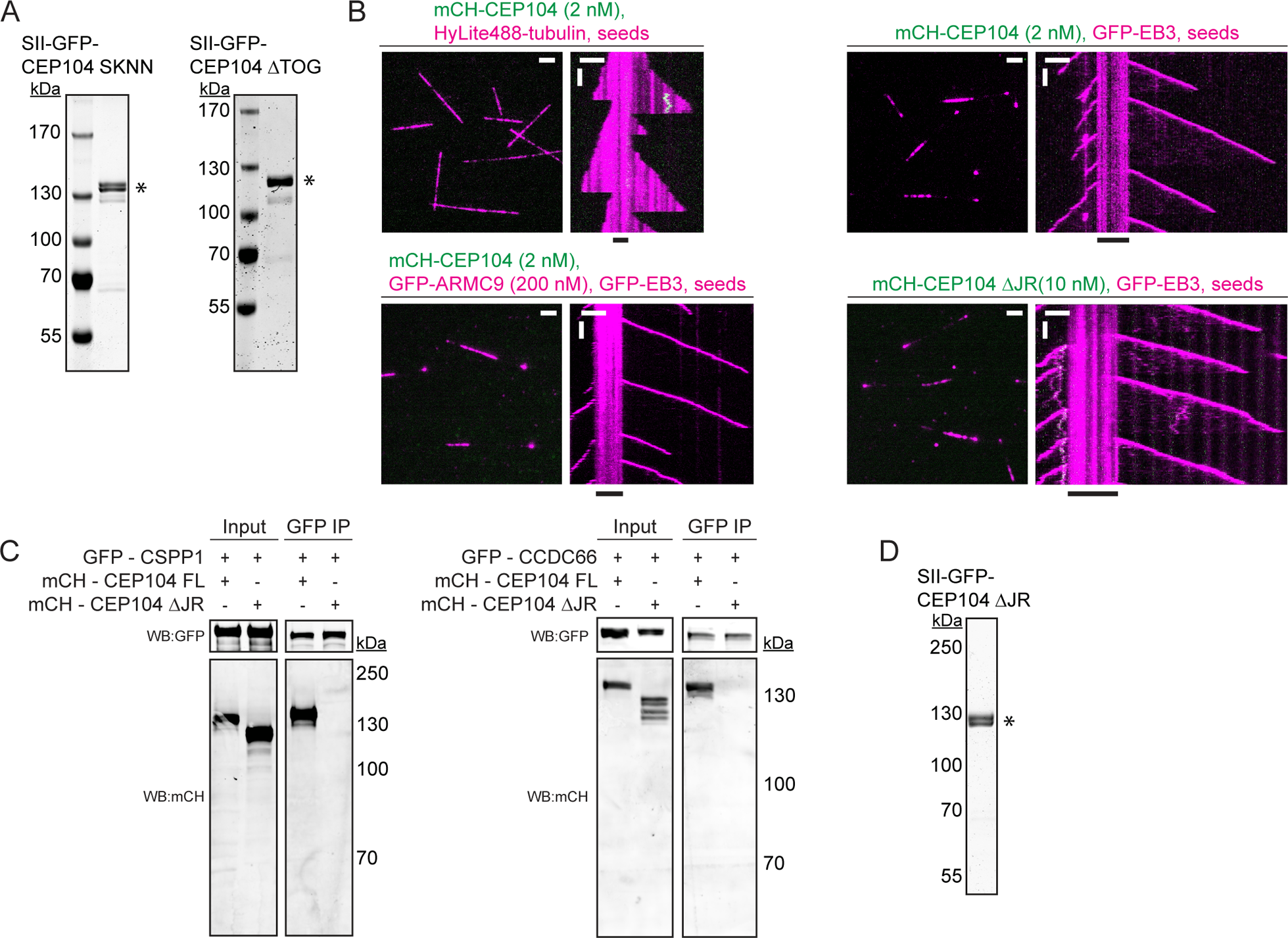
Characterization of CEP104 constructs *in vitro*. (A) Analysis of purified GFP-tagged CEP104 constructs by SDS-PAGE. Asterisks show protein bands. Protein concentrations were determined from a BSA standard (not shown). (B) Fields of view (left, scale bar 2 µm) and kymograph (right, scale bars 2 µm and 60 s) illustrating MT dynamics from GMPCPP-stabilized seed with either HiLyte-488-tubulin or GFP-EB3 and indicated concentrations and colors of CEP104 constructs. (C) Co-immunoprecipitation of either CSPP1 (left) or CCDC66 (right) with indicated CEP104 constructs, both CSPP1 and CCDC66 interact with the jelly-roll domain of CEP104. (D) Analysis of purified GFP-tagged CEP104 ΔJR construct by SDS-PAGE. Asterisk indicates the full-length protein band. Protein concentration was determined from a BSA standard (not shown).

**Figure S3 (Related to Figure 4).**
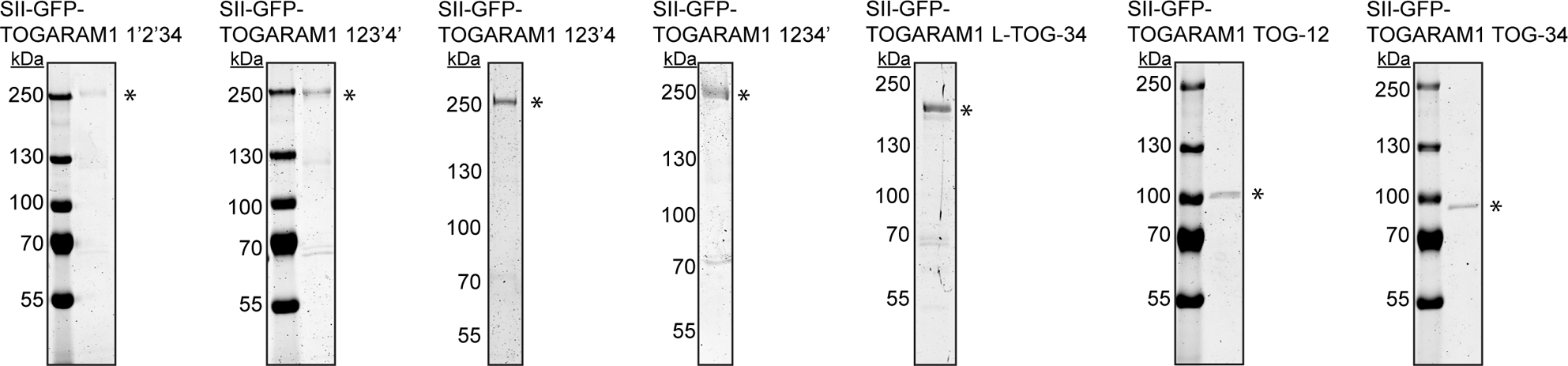
Characterization of TOGARAM1 constructs *in vitro*. Analysis of purified GFP-tagged TOGARAM1 constructs by SDS-PAGE. Asterisks show protein bands. Protein concentrations were determined from a BSA standard (not shown).

**Figure S4 (Related to Figure 5).**
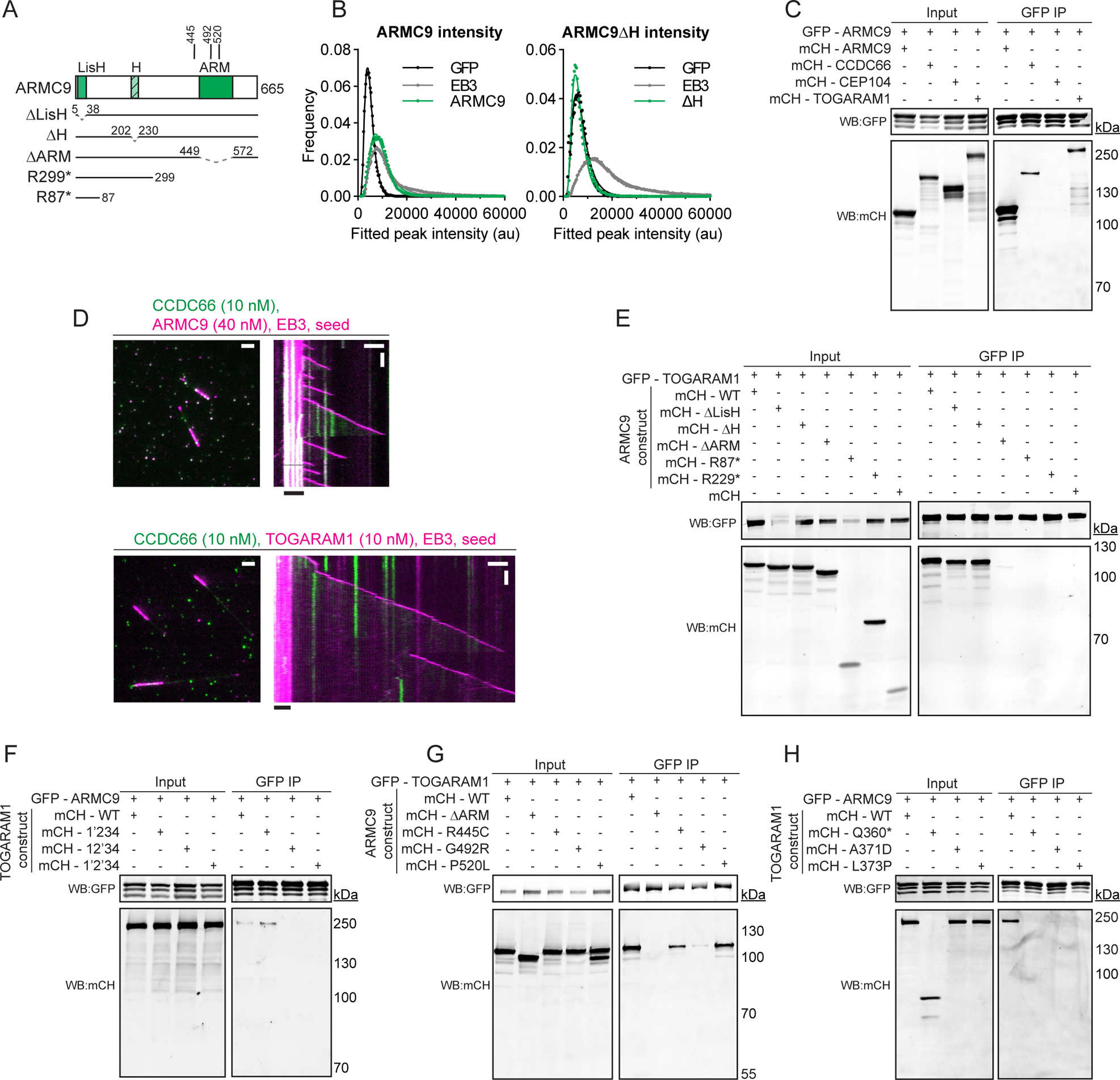
Mapping of ARMC9 and TOGARAM1 interaction. (A) Schematic representation of different ARMC9 constructs. (B) Histogram plots of fluorescent intensities of single GFP molecules, GFP-EB3 dimers, and full-length GFP-ARMC dimers (left) or GFP-ARMC9 ΔH molecules immobilized in separate chambers of the same coverslips. Number of molecules analyzed, left, right: GFP, n=18415, 14914; GFP-EB3, n= 132826, 163959; GFP-ARMC9 FL, n=99873; GFP-ARMC9 ΔH, 55479. (C) Co-immunoprecipitation of full-length ARMC9 with indicated proteins. (D) Fields of view (left, scale bar 2 µm) and kymograph (right, scale bars 2 µm and 60 s) illustrating MT dynamics from GMPCPP-stabilized seed with mCherry-EB3 and indicated concentrations and colors of ciliary tip module proteins. (E-F) Co-immunoprecipitation experiments of either full-length TOGARAM1 with indicated ARMC9 constructs (E) or full-length ARMC9 with indicated TOGARAM1 point mutations (F). ARMC9 and TOGARAM1 interact through ARM domain and TOG2 domain, respectively. (G-H) Co-immunoprecipitation experiments of either full-length TOGARAM1 with indicated ARMC9 Joubert syndrome mutations (G) or full-length ARMC9 with indicated TOGARAM1 Joubert syndrome mutations (H).

**Figure S5 (Related to Figure 5).**
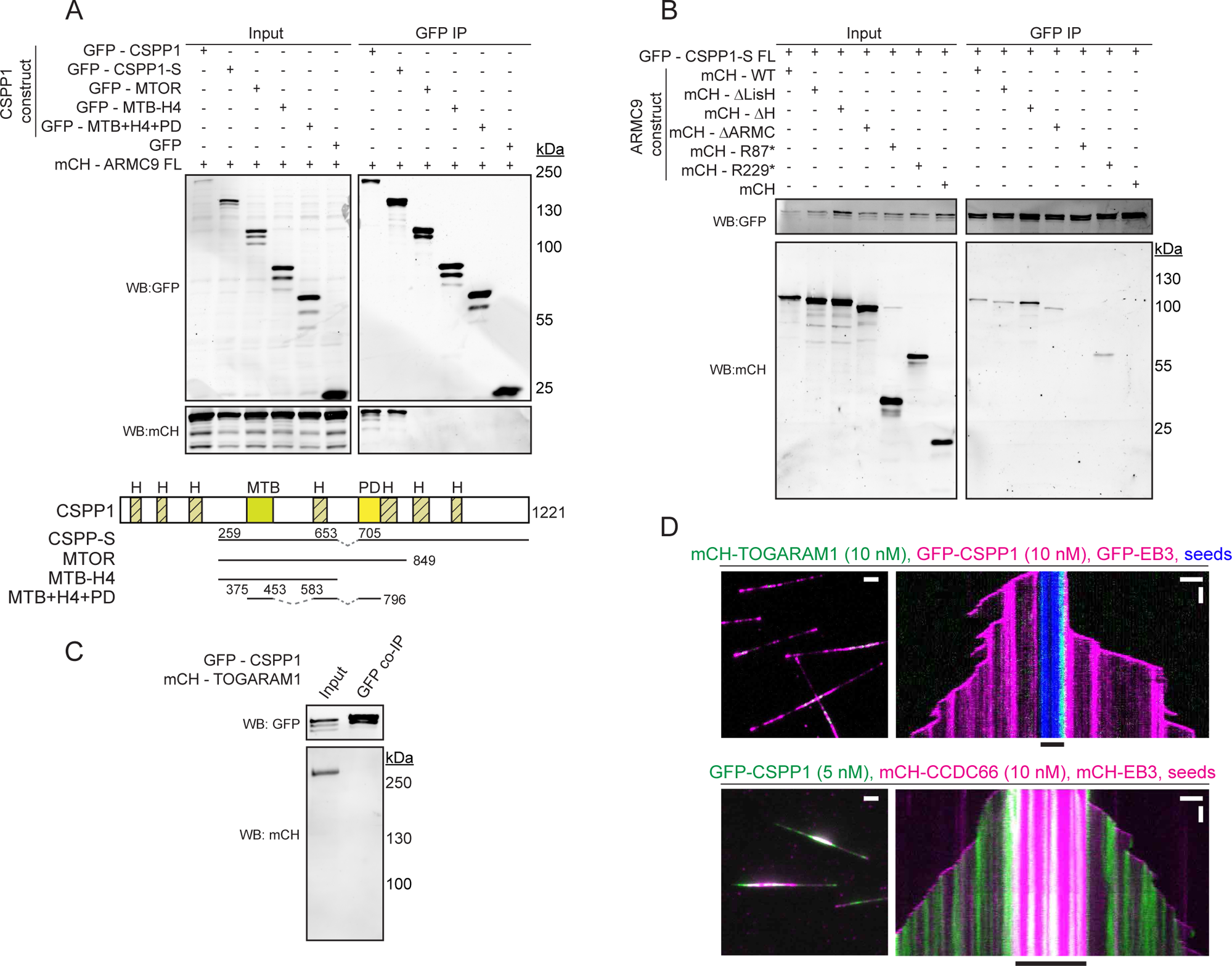
Mapping of ARMC9 and CSPP1 interaction. (A) Co-immunoprecipitation experiment of full-length ARMC9 with indicated CSPP1 constructs illustrated in schematic below. (B) Co-immunoprecipitation experiment of full-length CSPP1 with indicated ARMC9 constructs illustrated in Figure S4A. (C) Co-immunoprecipitation experiment of full-length CSPP1 with full-length TOGARAM1 shows no interaction. (D) Fields of view (left, scale bar 2 µm) and kymograph (right, scale bars 2 µm and 60 s) illustrating MT dynamics from GMPCPP-stabilized seed with GFP- or mCherry-EB3 and indicated concentrations and colors of ciliary tip module proteins.

**Figure S6 (Related to Figure 6).**
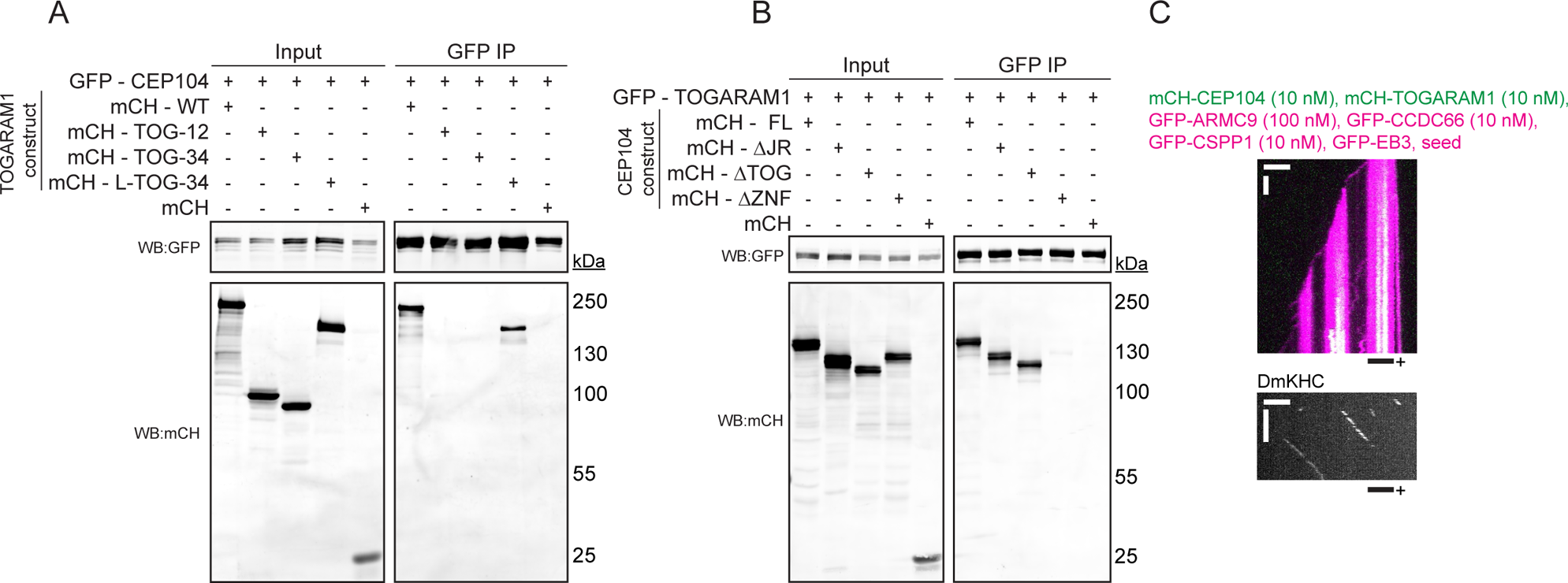
Mapping of CEP104 and TOGARAM1 interaction. (A-B) Co-immunoprecipitation of either wildtype CEP104 with indicated TOGARAM1 constructs (A) or wildtype TOGARAM1 with indicated CEP104 constructs (B), the two proteins interact between the linker of TOGARAM1 and the zinc finger of CEP104. (C) Kymographs illustrating mobility of DmKHC(1-421) on slow growing MT with all ciliary tip module proteins (top) and DmKHC (bottom) proving that the slow growing end of the MT is the plus end. Scale bars 2 µm and 60 s for both kymographs.

**Figure S7 (Related to Figure 7).**
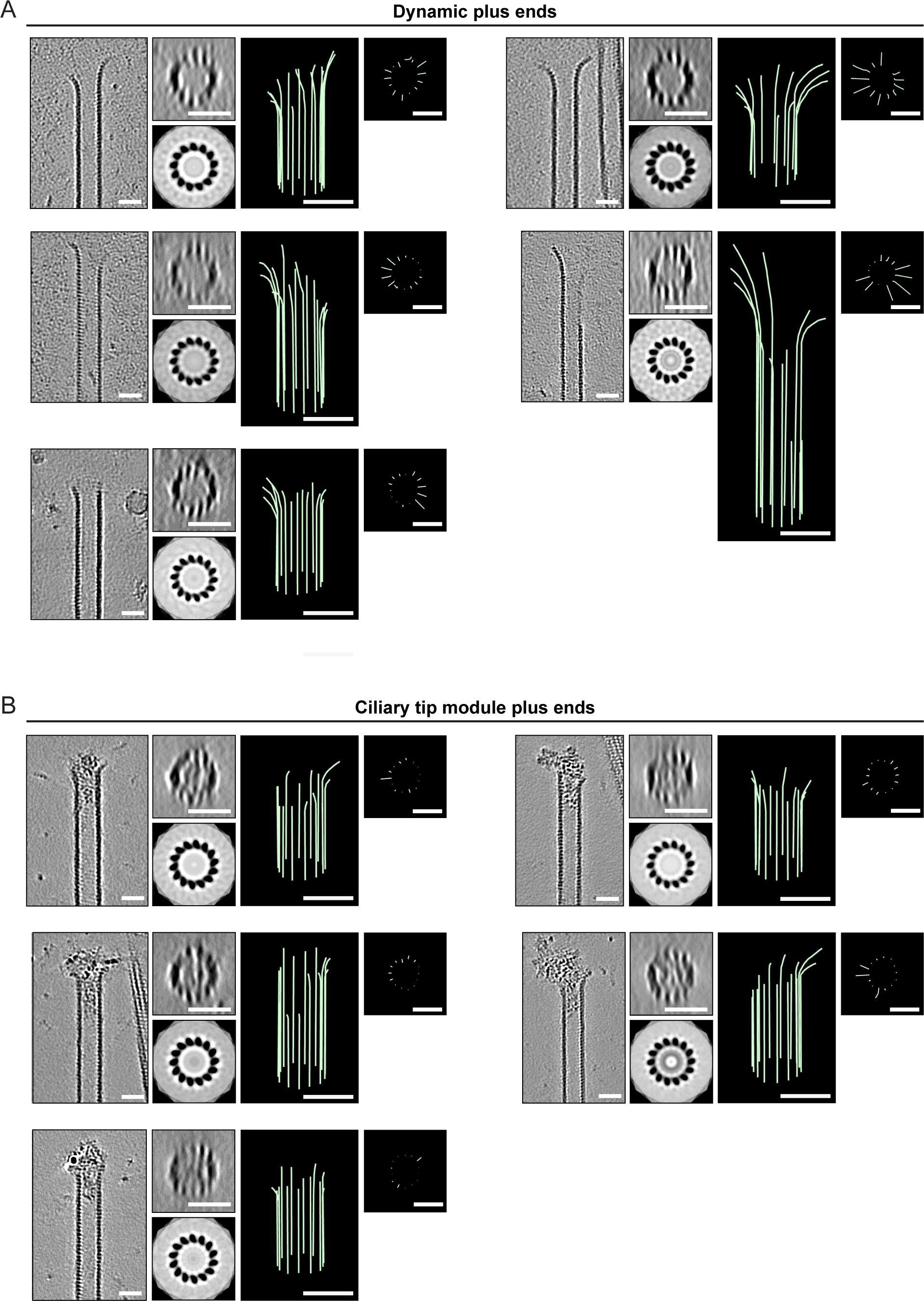
Characterization of MT plus ends by cryo-ET. (A, B) Representative examples of MT plus ends grown in presence of EB3 alone (A) or ciliary tip module and EB3 (B). Per MT, the following is shown from left to right; a slice of the denoised tomogram containing the MT plus end, an 8 nm thick transverse cross-section accompanied by rotational averaging analysis to determine MT polarity, and the corresponding 3D model of manual protofilament tracing accompanied by its transverse cross-section. Scale bars 25 nm.

## Notes

### Competing Interest Statement

The authors have declared no competing interest.

